# Multidrug-Resistant Avian Pathogenic *Escherichia coli* Strains and Association of Their Virulence Genes in Bangladesh

**DOI:** 10.1101/2020.06.30.180257

**Authors:** Otun Saha, M. Nazmul Hoque, Ovinu Kibria Islam, Md. Mizanur Rahaman, Munawar Sultana, M. Anwar Hossain

**Affiliations:** Department of Microbiology, University of Dhaka, Dhaka-1000, Bangladesh; Department of Gynecology, Obstetrics and Reproductive Health, Bangabandhu Sheikh Mujibur Rahman Agricultural University, Gazipur-1706, Bangladesh; Department of Microbiology, Jashore University of Science and Technology, Jashore-7408, Bangladesh

**Keywords:** Poultry, *Escherichia coli*, Phylotypes, Pathogenic, Antibiotic Resistance

## Abstract

The avian pathogenic *Escherichia coli* (APEC) strains are the chief etiology of avian colibacillosis worldwide. The present study investigated the circulating phylotypes, existence of virulence genes (VGs) and antimicrobial resistance (AMR) in 392 APEC isolates, obtained from 130 samples belonged to six farms using both phenotypic and PCR-based molecular approaches. Congo red binding (CRB) assay confirmed 174 APEC isolates which were segregated into 10, 9 and 8 distinctive genotypes by RAPD assay (discriminatory index, D=0.8707), BOX-PCR (D=0.8591) and ERIC-PCR (D=0.8371), respectively. The combination of three phylogenetic markers (*chuA, yjaA* and DNA fragment TspE4.C2) classified APEC isolates into B2_3_ (37.36%), A1 (33.91%), D2 (11.49%), B2_2_ (9.20%) and B1 (8.05%) phylotypes. Majority of the APEC isolates (75-100%) harbored VGs (*ial, fim*H, *crl, pap*C and *cjr*C), and of them, VGs (*pap*C and *cjr*C) and phylotypes (D2 and B2) of APEC had significant (p=0.004) association with colibacillosis. Phylogenetic analysis showed two distinct clades (Clade A and Clade B) of APEC where Clade A had 98-100.0% similarity with *E. coli* APEC O78 and *E. coli* EHEC strains, and Clade B had closest relationship with *E. coli* O169:H41 strain. Interestingly, phylogroups B2 and D2 were found in the APEC strains of both clades while the strains from phylogroups A1 and B1 were found in clade A only. In this study, 81.71% of the isolates were biofilm formers and possessed plasmids of varying ranges (1.0 to 54 kb). *In vitro* antibiogram profiling revealed that 100.0% isolates were resistant to ≥3 antibiotics, of which61.96%, 55.24, 53.85, 51.16 and 45.58 % isolates in phylotypes B1, D2, B2_2_, B2_3_ and A1, respectively were these antimicrobials. The resistance patterns varied among different phylotypes, notably in phylotype B2_2_ showing the highest resistance to ampicillin (90.91%), nalidixic acid (90.11%), tetracycline (83.72%) and nitrofurantoin (65.12%). Correspondence analysis also showed significant correlation of phylotypes with CRB (p=0.008), biofilm formation (p=0.02), drug resistance (p=0.03) and VGs (p=0.06). This report demonstrated that B2 and A1 phylotypes are dominantly circulating APEC phylotypes in Bangladesh; however, B2 and D2 are strongly associated with the pathogenicity. A high prevalence of antibiotic resistant APEC strains from different phylotypes suggest to use of organic antimicrobial compounds, and/or metals, and the rotational use of antibiotics in the poultry farms in Bangladesh.

## 1. Introduction

*Escherichia coli* is a ubiquitous a ubiquitous organism having the fabulous adaptive ability in diverse ecological niches including the intestine of animals and humans, and a pathogen that can induce enteric and extraintestinal infections (Jang et al., 2017; Solà-Ginés et al., 2015). In particular, avian pathogenic *E. coli* (APEC) is the main etiology of colibacillosis in poultry farms; a syndrome associated to diarrhea and/or enteritis, mild to severe septicemia, airsacculitis, perihepatitis, and pericarditis (Solà-Ginés et al., 2015). However, in most of cases, the fundamental cause of the disease remains unclear, since the infection with *E. coli* is associated to the presence of *Mycoplasma gallisepticum* or respiratory viruses, such as Newcastle virus or Infectious Bronchitis virus (Solà-Ginés et al., 2015). Moreover, the strains of *E. coli* also have zoonotic significance since they are known to cause infections both in humans and animals including birds (Kabiswa et al., 2018).

Currently, one of the greatest challenges for APEC is the difficulty of understanding their pathogenicity (Chakraborty et al., 2015). There are many approaches for molecular detection of various genomic fragments of the APEC, of them polymerase chain reaction (PCR) is one of the best approaches including studying their pathogenicity and/or virulence. The AFLP (amplified fragment length polymorphism) (Tikoo et al., 2001), RAPD (random amplified polymorphic DNA) (Yoon et al., 2016), repetitive sequence-based PCR genomic fingerprinting (Seurinck et al., 2003), multilocus sequence typing (MLST) (Moulin-Schouleur et al., 2007), BOX-PCR (Dombek et al., 2000), Clermont phylotyping (Clermont et al., 2013) and ERIC-PCR (Daga et al., 2019) are most widely used PCR-based techniques for identifying APEC strains. Among the mentioned techniques, MLST has discriminatory power and reproducibility for typing *E. coli* (Moulin-Schouleur et al., 2007), though it has some inherent limitations like relatively higher costs, unavailability and labor-intensiveness (Johnson et al., 2017; Salipante et al., 2015). On the contrary, there are high correlations between the Clermont phylogenetic grouping and MLST analysis (Gordon et al., 2008; Clermont et al., 2013). The *E. coli* strains can be grouped into different distinct phylogroups having originated from different ecological niches, and tendency to cause diseases through phylogenetic analysis using advanced molecular techniques (Pasquali et al., 2015; Aslam et al., 2014; Ghanbarpour et al, 2011; Wang et al., 2010; Dissanayake et al., 2008). Therefore, identifying and classifying the phylotype of an unknown strain can further expedite proper prevention and control programs, and also aid in designing rational treatment of infections caused by such strain (Müştak et al., 2015), because of its simplicity, quick and reproducible (Carli et al., 2015; Kabiswa et al., 2018).

The potentially pathogenic *E. coli* strains can be screened by different tests, like phenotypic assays of Congo red binding (CRB) (Knöbl, et al., 2011). Many researchers considered CRB assay as an epidemiological marker to identify the APEC strains (Fodor et al., 2010; Amer et al., 2015; Zahid et al., 2016). The binding of Congo red is associated with presence of virulence genes such as *omp*A, *iss, crl* and *fim*H and genes for multiple resistance to antibiotics (Fodor et al., 2010; Amer et al., 2015; Zahid et al., 2016). The functional amyloid fibers assembled by *E. coli* are called curli, and APEC are associated with curli production. As amyloid, curli fibers are protease resistant and bind to Congo red (CR) and other amyloid dyes (Stebbins et al., 1992; Reichhardt & Cegelski,2018). In several previous studies, a strong correlation between CRB and pathogenic properties of APEC isolates was also reported (Ahmad et al., 2009; Ezz El Deen et al., 2010; Amer et al., 2015; Zahid et al., 2016). The characteristic of CR binding constitutes a moderately stable, reproducible and easily distinguishable phenotypic marker, and thus, many researchers advocated the use of CR dye to distinguishing between pathogenic and non-pathogenic microorganisms in APEC study (Ahmad et al., 2009; Ezz El Deen et al., 2010; Amer et al., 2015; Zahid et al., 2016).

Several studies revealed the incidence of various phylotypes of APEC strains in combinations of virulence-associated genes (Ahmed and Shimamoto, 2013; Johnson et al., 2008; Moulin-Schouleur et al., 2007). These virulence factors are associated with various virulence genetic markers such as P fimbriae structural subunit (*papA*) and P fimbriae assembly (*papC*) (Johnson et al., 2003), *fim*A (encoding type 1 fimbriae), bundle-forming pilus (*bfp*) and aerobactin iron uptake system (*aer*) (López-Saucedo et al., 2003), *crl* (curli fimbriae) and many more which are linked to zoonotic concern (Knöbl et al., 2012). Among all the existent adhesion factors, the P fimbriae is one of the essential factors in pathogenesis of the poultry epithelial cells (Moulin-Schouleur et al., 2007). The P fimbriae are important factor for the beginning and expansion of human urinary tract infections (Campos et al., 2005); however, their role in the pathogenesis of avian have not yet clearly understood. The role of curli fimbriae which encodes for *crl* and *csg*A genes in the pathogenesis is poorly elucidated, though it facilitates the adherence of APEC strains to fibronectin and laminin (Amer et al., 2018; Borzi et al., 2018; Campos et al., 2005). Moreover, biofilm forming ability of the APEC strains is another vital virulence property (Kur, 2009) that justifies the reason for treatment failure using commercially available antimicrobials leading to persistence of the infections (Romling and Balsalobre, 2012). In addition, APEC isolates shared serotypes, virulence genes, and phylotypes with human dirrhoeagenic *E. coli* (DEC) isolates, which is a subsequent potential public health concern isolates (Xu et al., 2017; Ramadan et al., 2016).

Antibiotics resistance in bacteria is now a global public health concern, particularly this resistant property further aggravated due to indiscriminate use of antibiotics (Glass-Kaastra et al., 2014). The relation of antibiotics resistance especially in APEC phylotypes had previously been discussed (Borzi et al., 2018; Mahmoudi-Aznaveh et al., 2013). There are reports that resistant *E. coli* strains might enhance antimicrobial resistance in other organisms (pathogenic and nonpathogenic) within gastrointestinal tract of the chicken (Tello et al., 2012), and help in transmitting and disseminating drug-resistant strains and genes between animal and human pathogens (Kabiswa et al., 2018; Tello et al., 2012).

Many of the earlier studies have explore the association between pathogenic traits of APEC and their virulence gene repertoire (Azad et al., 2019; Sarker et al., 2019; Logue et al., 2017; Reza et al., 2009); however, potential pathogenic traits associated with phylotypes warrants attention into the potential role of virulence genes (VGs), MDR properties, and their cross-talk with specific phylotypes. This study was therefore aimed to investigate the phylotyping, potential VGs, and antimicrobial resistance (AMR) in APEC isolates. Furthermore, we also studied the role of phylotyping in APEC to be associated with VGs carriage and AMR in infected poultry samples of Bangladesh.

## 2. Materials and Methods

### 2.1 Sample collection and processing

We collected 130 samples (healthy: 35 & diseases: 95) including droppings (n=30), cloacal swabs (n=27), poultry feed (n=11), handler’s swab (n=9), egg surface swab (n=10), feeding water (n=10) and internal organ (liver, n=33) from 6 commercial poultry farms belonged to Narsingdi (23.9193° N, 90.7176° E), Narayangonj (3.6238° N, 90.5000° E), and Manikgonj (23.8617° N, 90.0003° E) districts of Bangladesh during April 2017 to March 2018 (Supplementary Table 2). Diseased chickens (colibacillosis) were confirmed by observing the clinical syndrome associated to diarrhea and/or enteritis, mild to severe septicemia, airsacculitis, perihepatitis, and pericarditis (Solà-Ginés et al., 2015). Collected samples were put into sterile plastic bags, carefully labeled, packed, cooled in icebox, transported subsequently to the laboratory, and stored at 4°C. Further processing of all samples was done for microbiological analyses (Osman et al., 2018).

### 2.2 Isolation and identification of pathogenic *E. coli*

We adopted the method of Knobl et al. (2012), and modified protocol of Food and Drug Administration Bacteriological Analytical Manual (FDA-BAM) guidelines for isolating and identifying APEC (Feng et al., 2015). In brief, loopful inoculums from each sample was inoculated into previously prepared nutrient broth (NB) and incubated for 24 h at 37°C. A small amount of inoculum from NB was streaked onto MacConkey (MC) agar and Eosin Methylene Blue (EMB) agar for 24 h at 37°C for selective growth. The colonies showing characteristic metallic sheens on selective Eosin methylene blue (EMB) agar (3–5 colonies from each sample) were selected to further biochemical tests (indole, methyl-red, catalase, citrate and Voges– Proskauer) for confirmatory identification of *E. coli* (Osman et al., 2018).

### 2.3 Phenotypic virulence assays

The assays for virulence properties of *E. coli* were performed according to the (FDA-BAM guidelines.

#### 2.3.1 Congo red binding assay

The Congo red binding (CRB) ability of the APEC isolates was determined using agar plates supplemented with 0.003% CR dye (Sigma, USA) and 0.15% bile salts. Bacterial suspension (5 µL) was streaked onto the plates, and the plates were incubated at 37°C for 24 h. The strong biofilm-producing isolates were visualized as deep brick red-colored. For more confirmation, colonies were also consecutively examined following 48 and 72 h of incubation and the results were interpreted as +++, ++ and + depending on their color intensity (Reichhardt and Cegelski, 2018; Bist et al., 2014).

#### 2.3.2 Production of hemolysins and swimming motility assays

The *E. coli* hemolytic activity of the CRB positive isolates was evaluated by streaking blood agar base plates with 5% sheep blood. After 24 h incubation at 37°C, plates were examined for signs of β-hemolysis (clearing zones around colonies), α-hemolysis (a green-hued zone around colonies) or γ-hemolysis (no halo around colonies) (Maragkoudakis et al., 2006). We performed motility assay by following the previously described protocols (Wang et al., 2016). In brief, bacterial cultures were stabbed onto motility indole urease (MIU) soft agar motility tubes (0.5% agar), and the tubes were incubated at 37°C. Finally, we measured the bacterial motility haloes after 24 and 48 h of incubation (Wang et al., 2016).

#### 2.3.3 Biofilm formation assay

The biofilm formation assay of randomly selected 82 CRB positive *E. coli* isolates was performed by using 96 well microtiter plate methods to quantitatively measure the attachment, and biofilm formation on solid surfaces (plastic) during static conditions (Murray et al., 2017). The assay was performed in duplicate in the 96 well tissue culture plates. The observed optical density (OD) was evaluated to determine the biofilm-forming ability of the isolates on a 4-grade scale (non-adherent, weakly adherent, moderately adherent and strongly adherent). This 4-grade was determined by comparing OD with cut-off OD (ODc) (three standard deviation values above the mean optical density of the negative control). Furthermore, the biofilm surface after 24 h was stained with biofilm viability kit to observe the proportion of live or active cells (fluorescent green) and dead or inactive cells (fluorescent red) using fluorescence microscope and DP73 digital camera (40X objective). After that the microscopic images were analyzed by the image J software (Murray et al., 2017).

### 2.4 DNA extraction

Genomic DNA of *E. coli* was extracted from overnight culture by the boiled DNA extraction method (Sultana et al., 2018). Briefly, the samples were centrifuged at 15,000g for 15 min (Sultana et al., 2018). We eliminated the supernatant, resuspended the pellet in molecular biology-grade water, and centrifuged at 15,000 g for 10 min. Again, the supernatant was eliminated, the pellet was resuspended in 40 µl of molecular biology grade water, subjected to boiling at 100°C in a water bath for 10 min, cooled on ice and centrifuged at 15,000g for 10 s (Hossain et al., 2018). The extracted DNA was quantified using a NanoDrop ND-2000 spectrophotometer.

### 2.5 Molecular typing methods

Species identification of *E. coli* in 174 CRB positive isolates were confirmed using *E. coli* specific PCR such as random amplification of polymorphic DNA (RAPD) (Yoon et al., 2016), box elements (BOX-PCR) (Dombek et al., 2000) and enterobacterial repetitive intergenic consensus (ERIC-PCR) (Daga et al., 2019). The optimized protocol for RAPD PCR was performed by using (5’-GCGATCCCCA-3’) primer (Hossain et al., 2018), while the ERIC-PCR was done with ERIC1R (5′-ATGTAAGCTCCTGGGGATTCAC-3′) and ERIC2 (5′-AAGTAAGTGACTGGGGTGAGCG-3′) primers (Daga et al., 2019) and the BOXA1R (5′-CTACGGCAAGGCGACGCTGACG-3′) primer was used for BOX PCR (Dombek et al., 2000).

### 2.6 Clermont’s phylogenetic typing

The isolates were subjected to a phylotyping group using the Clermont et al. method (Clermont et al., 2013). The reaction mixture (15 μl) contained about 7.5 μL of GoTaq® G2 Green Master Mix (Promega, USA), 1.0 μL of primer, 4.5μL nuclease-free water (Promega, USA), and 1.0 μL of template genomic DNA. Cycling conditions were as follows: 5 min initial denaturation at 94°C, followed by 30 cycles of 5 s at 94°C, 25 s at 59°C; with a final extension of 5 min at 72°C. PCR products were separated and visualized on 1.2% agarose gels in 1-Trisborate-EDTA (TBE) buffer using ethidium bromide staining. Following electrophoresis, the gel was photographed under UV light, and the strains were assigned to the phylotypes B2 (*chu*A+, *yja*A+), D (*chu*A+, *yja*A-), B1 (c*hu*A-, TspE4.C2+), C (*chu*A-, *yja*A+, TspE4.C2-), E (*chu*A+, *yj*aA-, TspE4.C2+), F (*chu*A+, *yja*A-, TspE4.C2-), or A (*chu*A-, TspE4.C2-). For better strain-level discrimination, the subgroups or phylotypes were determined as follows: subgroup A0 (group A), *chu*A-, *yja*A-, TspE4.C2-; subgroup A1 (group A), *chu*A-, *yja*A+ TspE4.C2-; group B1, *chu*A-, *yja*A-, TspE4.C2+; subgroup B2_2_ (group B2), *chu*A+, *yja*A+, TspE4.C2-; subgroup B2_3_ (group B2), *chu*A+, *yja*A+, TspE4.C2+; subgroup D1 (group D), *chu*A+, *yja*A-, TspE4.C2- and subgroup D2 (group D), *chu*A+, *yja*A-, TspE4.C2+ (Iranpour et al., 2015).

### 2.7 Sequencing and phylogenetic analysis based on 16S rRNA gene Sequence

*E. coli* isolates from each phylotype along with others typing (RAPD, ERIC and BOX-PCR) were selected for 16S rRNA gene PCR using universal primers (27F 5’-AGAGTTTGATCMTGGCTCAG-3’ and 1492R 5’-TACGGYTACCTTGTTACGACTT-3’) followed by sequencing at First Base Laboratories SdnBhd (Malaysia) using Applied Biosystems highest capacity-based genetic analyzer (ABI PRISM^®^ 377 DNA Sequencer) platforms with the BigDye^®^ Terminator v3.1 cycle sequencing kit chemistry (Masomian et al., 2016). Initial quality controls of the generated raw sequences were performed using SeqMan software and were aligned with relevant reference sequences retrieved from NCBI database using Molecular Evolutionary Genetics Analysis (MEGA) version 7.0 for bigger datasets (Kumar et al., 2016). Evolutionary distances were computed using the Kimura-Nei method and the phylogenetic tree was constructed by applying the neighbor-joining method (Saitou, 1987). The percentage of replicate trees in which the associated taxa clustered together in the bootstrap test (1000 replicates) is shown next to the branches. The evolutionary distances were computed using the Kimura 2-parameter method and is in the units of the number of base substitutions per site.

### 2.8 Antimicrobial sensitivity testing

Kirby-Bauer disc diffusion method on Mueller-Hinton agar was used for antimicrobial susceptibility testing of various *E. coli* pathotypes according to the guidelines of the Clinical and Laboratory Standards Institute (Abbey and Deak, 2019). Antibiotics were selected for susceptibility testing corresponding to a panel of antimicrobial agents of interest to the poultry industry and public health in Bangladesh. Randomly selected representative 114 *E. coli* isolates from all phylotypes were evaluated for antimicrobial susceptibility to 13 antibiotic discs belongs to 11 different antimicrobial classes including penicillins (ampicillin,10 µg); tetracyclines (doxycycline,30 µg; tetracycline,30µg); nitrofurans (nitrofurantoin,300 µg); lipopeptides (polymexin B,30µg); monobactams (aztreonam, 30µg); quinolones (ciprofloxacin, 10µg; nalidixic acid, 30µg); cephalosporins (cefoxitin, 30µg), penems (imipenem,10µg); aminoglycosides (gentamycin, 10µg); phenols (chloramphenicol, 30 µg); sulfonamide (sulfonamide,250µg) and macrolides (azithromycin, 15µg) (Oxoid, UK). Finally, the findings were recorded as susceptible, intermediate and resistant according to Clinical and Laboratory Standards Institute (Abbey and Deak, 2019) break points.

### 2.9 Detection of virulence genes

We surveyed 13 virulence genes (VGs) that are normally studied in APEC strains. The selected genes included: *eae* (Intimin) (López-Saucedo et al.,2003), *stx*1 (shiga toxins 1) (López-Saucedo et al.,2003), *stx*2 (shiga toxins 2) (López-Saucedo et al.,2003, *fim*H (type 1 fimbriae) (Hojati et al., 2015; Borzi et al., 2018), *hly*A (hemolysin) (Srivani et al., 2017; Borzi et al., 2018), *papC* (P fimbriae) (Yamamoto et al., 1995; Borzi et al., 2018), *lt* (heat-labile enterotoxin) (Houser et al., 2008), *bfp*A (bundle-forming pilus) (López-Saucedo et al.,2003), *crl* (curli fimbriae) (Knöblet al., 2012; Reichhardt et al., 2015; Amer et al., 2018; Reichhardt et al., 2018), *uid*A (β-d-glucuronidase) (Tsai et al.,1993), *agg*R (aggregative adherence regulator) (Toma et al., 2003), *ial* (invasion-associated locus) (López-Saucedo et al., 2003) and *cjr*C (putative siderophore receptor) (Mao et al., 2012). Each PCR reaction contained 2 *µ*L DNA template (300 ng/*µ*L), 10 *µ*L PCR master mix 2X (Go Taq Colorless Master Mix) and 1*µ*L (100 pmol/*µ*L) of each primer in each tube. The PCR amplifications were conducted in thermo cycler and the cycling conditions were identical for all the samples as follows: 94 °C for 5 min; 35 cycles of 1min at 94 °C, 1 min at 50-60 °C, and 1 min at 72 °C; and 72 °C for 7 min. PCR amplicons were visualized on 1.5% agarose gel prepared in 1× TAE buffer. After gel electrophoresis, the images were captured using Image ChemiDoc™ Imaging System (Bio-Rad, USA) (Knöbl et al., 2012; Mao et al., 2012).

### 2.10 Plasmid DNA isolation

The plasmid DNA of *E. coli* isolates was isolated following the method previously reported by different researchers (Rakhi et al., 2019; Hoque et al., 2018). Briefly, plasmid DNA of 45 randomly selected *E. coli* isolates was extracted using Wizard® Plus SV Mini preps plasmid DNA Purification kit (Promega, USA) according to manufacturer’s instruction and was electrophoresed in 0.8% agarose gel (Rakhi et al., 2019). *E. coli* V517 strain was used in the present study for molecular weight determination as control.

### 2.11 Discriminatory index (D)

The discriminatory index (D) was measured for each sample by entering data onto the following site: http://insilico.ehu.es/mini_tools/discriminatory_power/index.php (Bikandi et al., 2004).

### 2.12 Correspondence analysis

Correspondence analysis (CA) was used to study and compare the categories of molecular typing (phylotypes), the origin of sample groups (poultry feed and water, handler, egg surface, droppings, cloacal swab, liver), and pathogenic intensities (pathogenic genes, biofilm formation and drug sensitivity) using Pearson correlation tests through the IBM SPSS Statistics 20.0 package. A two-dimensional graph was used to show the relationship between the categories in CA analysis where the value of the third dimension is shown in parenthesis (Coura et al., 2017). The association of different sample types with molecular typing and pathogenic intensities were represented using the circular plot. The plot was visualized using OmicCircos (Hoque et al., 2020).

### 2.13 Statistical analysis

We used the SPSS software for Windows, version 20.0 (SPSS Inc., Chicago, IL, United States) for statistical analysis (Hoque et al., 2019). Comparison among frequencies of occurrence of each phenotypic or genotypic feature in *E. coli* isolates from poultry originated samples were carried out by contingency table χ2 tests (at p<0.05). To analyze motility, hemolysin and biofilm assay data we applied one-way analysis of variance (ANOVA), and two-way ANOVA was performed to analyze the typing methods and antimicrobial susceptibility tests. The association between molecular typing and sample types, and molecular typing and pathogenic intensities were calculated using the chi-square test. The result was considered to be significant at p≤0.05.

## 3. Results

### 3.1 Phenotypic characterization of APEC: isolation and identification

A total of 130 poultry samples (droppings, 30; cloacal swabs, 27; feeds, 11; handler’s swab, 9; egg surface swab, 10; feeding water, 10; liver, 33) were screened for phenotypic identification of *E. coli*. According to the microbiological analysis of the samples, 392 isolates were obtained through selective identification in EMB and MacConkey agar (metallic sheen on EMB agar plates and pink colonies on MacConkey agar) and biochemical tests followed by Congo red binding (CRB) assay. Results from the CRB assay revealed that 44.39% (174/392) of the isolates were avian-pathogenic *E. coli* (APEC), of which 24.71% (43/174) isolates were from healthy birds and 75.29% (131/392) from diseased birds. Of these APEC isolates, 46.55% (81/174) were retrieved from Narayangonj district followed by 39.38% and 32.69%, respectively from Manikgonj and Narsigdi districts. The distribution of pathogenic *E. coli* in different samples have been represented in Figure 1, and among them 33.33% *E. coli* were isolated from droppings, followed by liver (19.54%), cloacal swab (17.82%), handler swab (10.34%), feeding water (9.20%), feeds (5.17%) and egg surface swabs (4.60%), and (Supplementary Table 2). The frequency of the detected bacterial isolates also significantly (p=0.019) varied within the three sampling sites.

**FIGURE 1.**
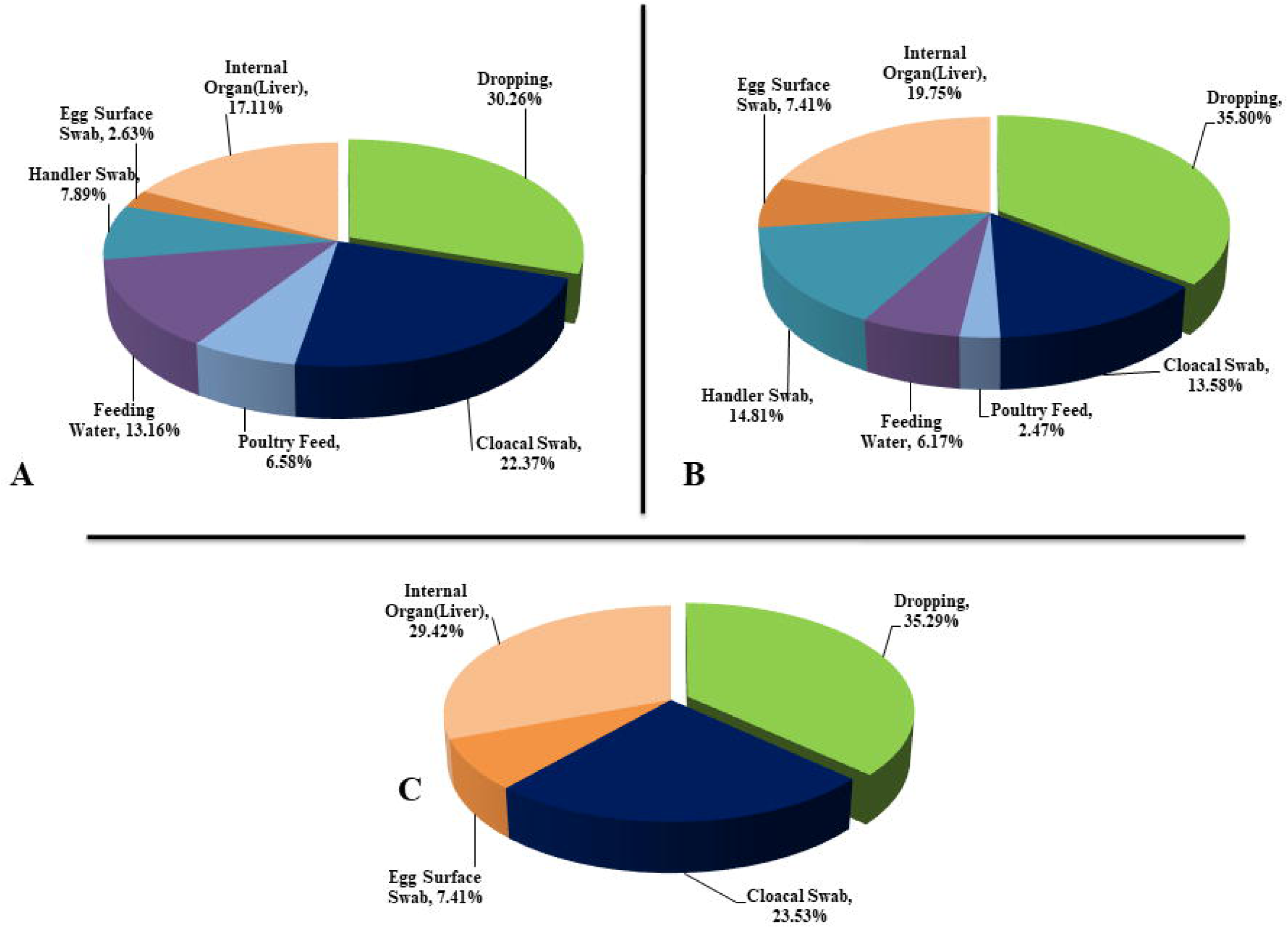
Occurrence of *Escherichia coli* in different types of poultry samples. Microorganisms identified in A (Dhamrai, Manikgonj); B (Rupganj, Narayangonj); and C (Monohardi, Narshingdi)) poultry samples. The Dhamrai, Rupganj and Monohardi region includes 76, 81 and 17 isolates, respectively.

### 3.2 Genetic diversity of APCE by molecular fingerprinting

In this study we used RAPD, ERIC and BOX PCR to analyze the genetic diversity and relatedness in 174 isolates of *E. coli* originated from different poultry samples. RAPD showed 10 different patterns among the isolates and the reproducibility of the RAPD technique was analyzed by repeated testing. (Supplementary Figure 1). In case of ERIC and BOX PCR, the number of DNA bands for different *E. coli* isolates were 1-5 and 1-7, respectively and thus, the isolates were differentiated into 8 and 9 groups through ERIC and BOX PCR, respectively (Supplementary Figures 2, 3). Therefore, RAPD fingerprint of DNA showed the highest genetic diversity among the isolates followed by BOX and ERIC PCR. The discriminatory indices (D) for RAPD, ERIC and BOX PCR were 0.8707, 0.8371 and 0.8591, respectively for all isolates. The diversity of the isolates was also measured through the principle component analysis (PCA). The PCA results revealed that the genetic diversity of *E. coli* isolates did not vary significantly according to molecular typing systems since in all typing methods group 1-4 clustered in the same quadrant of the PCA plot (Figure 2).

**FIGURE 2.**
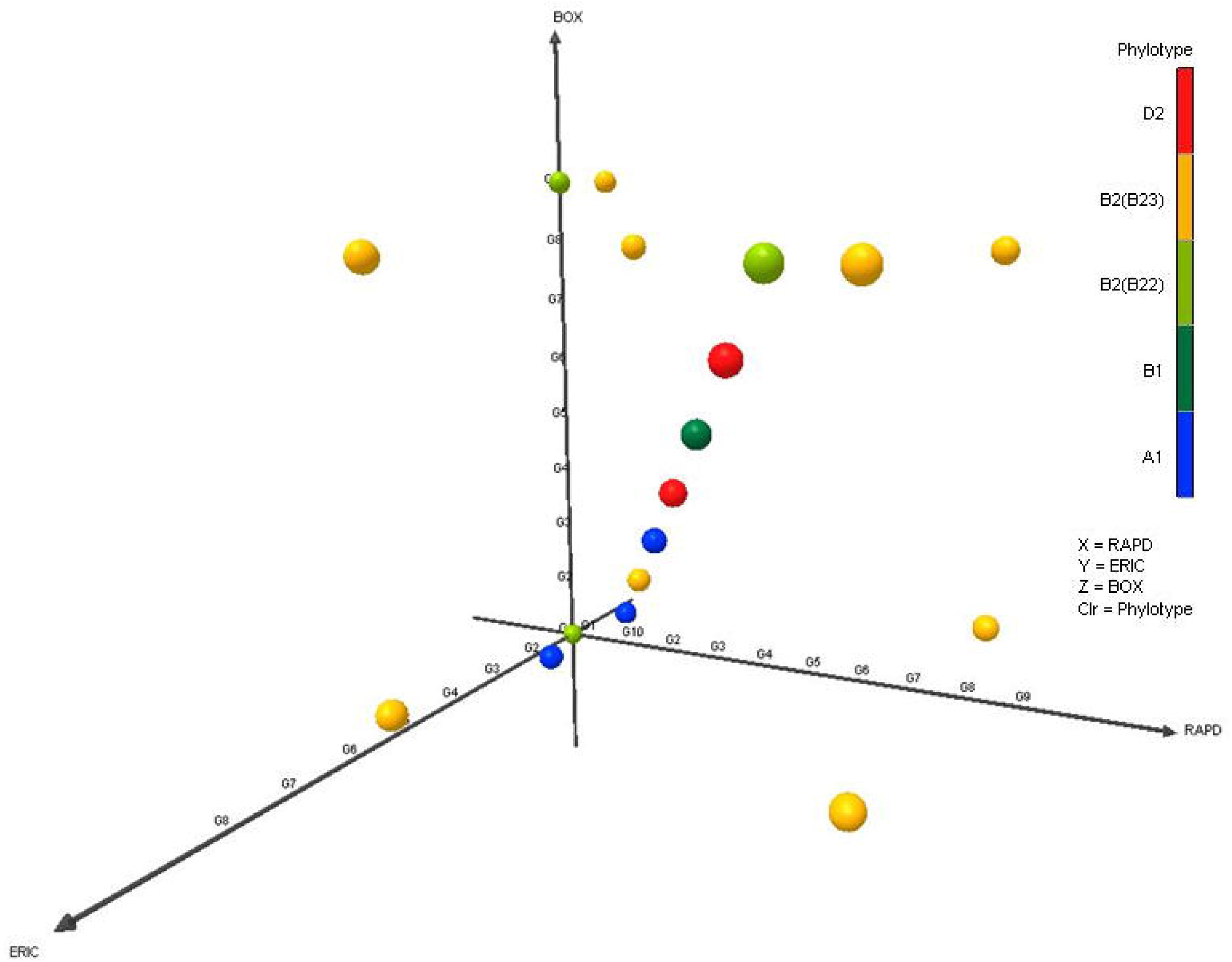
The diversity of APEC isolates according to various typing systems. The Principle Components Analysis (PCA) plots represent the distribution of different phylotypes. Orange is for phylotype D2, blue is for A1, Yellow is for B2_3_, Green is for B2_2_ and dark red green is for B1.

### 3.3 Phylogenetic distribution of APEC in poultry isolates

The distribution of 174 *E. coli* isolates belonged to five phylotypes (A_1_, B1, B2_2_, B2_3_ and D2). However, any gene combination for Phylotype E, C, and F were absent in the APEC isolates of the current study (Supplementary Table 2; Supplementary Figure 5). In the comparative analysis, we found that majority of poultry APEC isolates were affiliated to phylotype B2_3_ comprising 37.36% (65/174) followed by A1 (33.91%), D2 (11.49 %), B2_2_ (9.20 %) and B1 (8.05 %) (Figure 3, Supplementary Table 2). In this study, phylotype A1 were more prevalent (55.81%) in the samples from healthy birds while phylotype B2_3_ (41.22%) were more prevalent in the samples of infected birds followed by A1 (26.72%), D2 (14.50%), B2_2_ (11.45%) and B1 (6.11%) (Supplementary Table 2). Our results demonstrated that *E. coli* phylotypes distribution differed significantly (p=0.002) across the study areas. The isolates from Manikgonj district were segregated into phylotype B2_3_ (36.84%), A1 (30.26%), B1 (14.47%), D2 (11.84%) and B2_2_ (6.58%) while those from Narayangonj district were segregated into phylotypes A1 and B2_3_ (37.03%, each), B2_2_ (12.35%), D2 (9.99%) and B1 (3.70%). Conversely, none of the isolates from Narsingdi district harbored phylotype B1. In this study, the phylotype B2 (B2_2_ and B2_3_) and A1 were more prevalent in all of the *E. coli* isolates, however, B1 and D2 phylotypes were not found among the isolates of poultry feed and egg surface swab (Supplementary Table 2).

**FIGURE 3.**
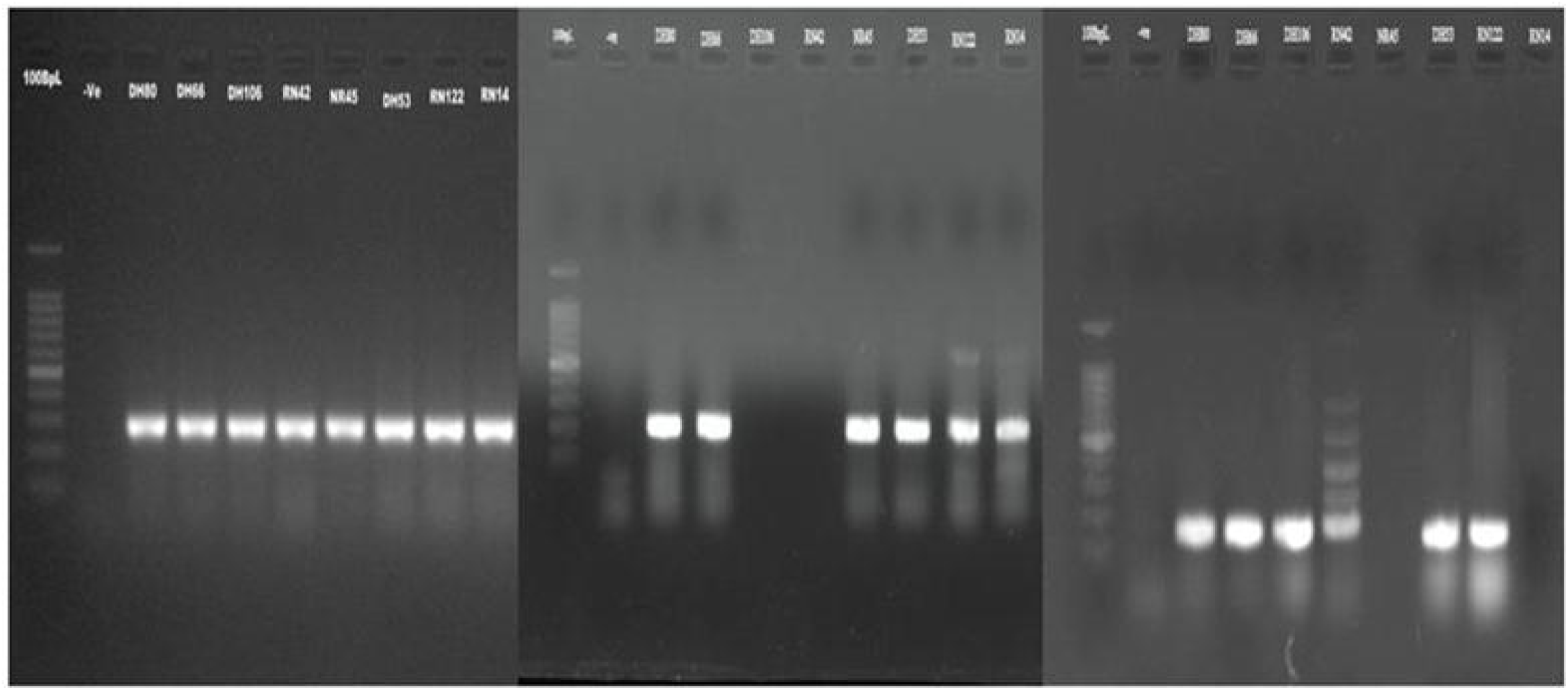
Phylotypes of APEC isolates detected by PCR method. A) (*chu*A: 279bp); B) (*yja*A: 211bp); C) (tspE4.C2:152bp); Here, Lane 1 is molecular ladders (100bp), Lane 2 is negative blank control and Lanes 3–10 are the strains DH80, DH66, DH106, RN42, NR45, DH53, RN122, RN14 respectively. DNA bands at the appropriate position was observed in *E. coli* strains DH66, DH80, DH53, RN122 denoted as phylotype B2(B2_3_) (*chu*A+, *yja*A+, tspE4.C2+); *E. coli* strain DH106, RN42 denoted as phylotype D2 (*chu*A+, *yja*A-, tspE4.C2+); *E. coli* strain NR45, RN14 denoted as phylotype B2(B2_2_) (*chu*A+, *yja*A+, tspE4.C2-).

### 3.4 Biofilm formation (BF) assay

As quantified in crystal violet assay, BF bacteria were divided into four groups based upon OD600 of the bacterial biofilm: non-biofilm forming (NBF), weak biofilm forming (WBF), moderate biofilm forming (MBF) and strong biofilm forming (SBF) bacteria. In this study, the average OD of the negative control was 0.028±0.002 and the cutoff OD value was set as 0.045. The isolates which have OD value ≤0.045 were considered as NBF. We found that 81.71% (67/82) of the *E. coli* isolates were BF representing 11(61.11%) and 56 (87.5%) isolates from healthy and diseased birds, respectively (Supplementary Table 2). By comparing the category of BF, we demonstrated that 30.49%, 26.83%, 24.39% and 18.29% isolates were SBF, MBF, WBF and NBF, respectively (Figure 4A, Supplementary Table 2). Of the SBF isolates, 4 (5.97%) and 21(31.34%) were found in healthy and infected birds, respectively (Supplementary Table 2). Microscopic observation followed by 3D image analysis revealed that the intensity of green fluorescence remained higher indicating that a large number of cells were viable and attached to the surface (Figure 4B). The isolates having SBF properties belonged to pathogenic *E. coli* (B2_2_, 42.86; B2_3_, 29.73; D2, 54.55) and thus, D2 isolates of pathogenic *E. coli* had the highest (54.55%) SBF ability (Figure 4A). Notably, all of the isolates from phylotype D2 had BF ability followed by phylotype B2 (86.36%). Interestingly, in both healthy and diseased birds phylotype B2_3_ showed the highest BF ability (Supplementary Table 2).

**FIGURE 4.**
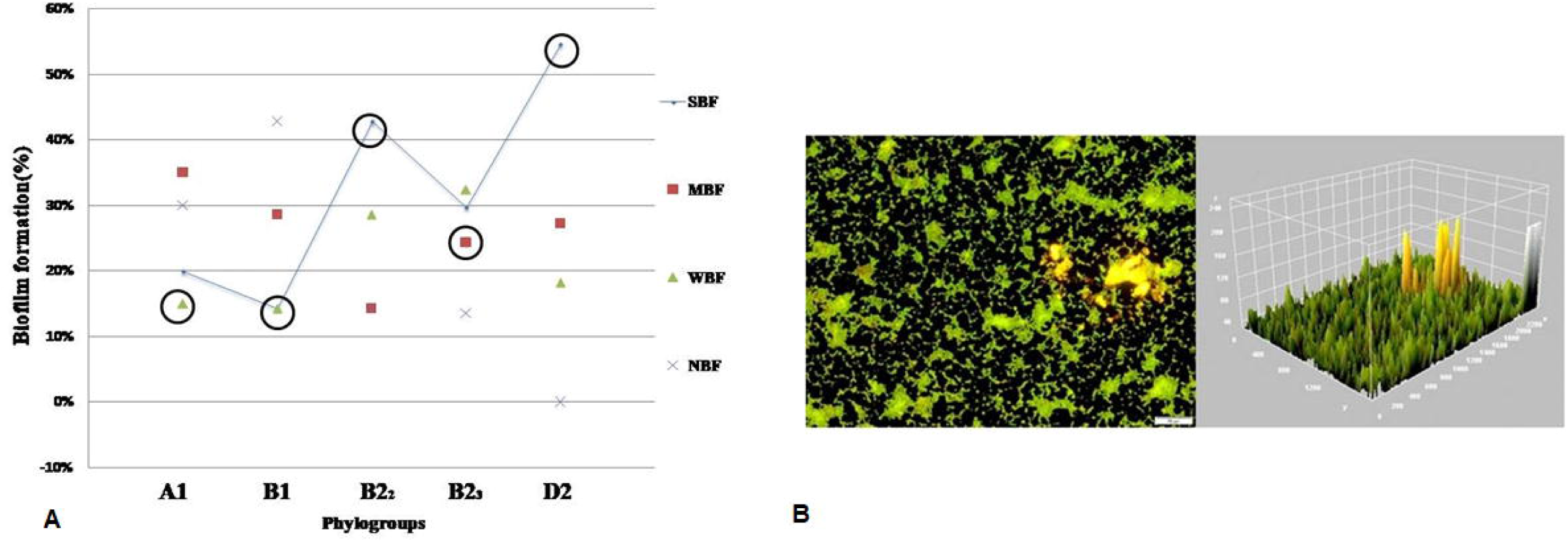
The biofilm formation ability of APEC isolates. A) Diagrammatic representation of Biofilm formation of various phylogroups. Here the categories of biofilm formation are strong biofilm formers (SBF), moderate biofilm formers (MBF), weak biofilm formers (WBF) and non-biofilm formers (NBF). Solid line with circle represented the SBF ability fluctuation between the phylotypes. X axis represent the phylogroups. Y axis represented the zone of percentages of biofilm forming isolates. B) Fluoroscence microscopy images of isolate (RN3 (2)) under 20x magnification. Biofilm stained with Film tracer LIVE/DEAD biofilm viability kit. Live or active cells are fluorescent green while the dead or inactive cells are fluorescent red. Surface plot of three dimensional (3D) volume image (center image) and cross section of 3D volume image (right side image) show the distribution of live and dead cells throughout biofilm layers.

### 3.5 Distribution of virulence genes of *E. coli* in poultry isolates

The possible association of different virulent genes (VGs) was screened through PCR in 123 APEC isolates according to their phylogroups. In this study, thirteen probable APEC associated VGs including the diarrheagenic and septicemic genes encoding for *eae, stx*1, *stx*2, *fim*H, *hly*A, *pap*C, *lt, bfp*A, *crl, uid*A, *agg*R, *ial* and *cjr*C were screened. The virulence genotyping showed that none of the APEC isolates harbored genes coding for *eae, stx1, stx2, hly*A, *bfp*A and *fli*C (Supplementary Figure 4). The distribution of the six apparently higher prevalent VGs such as *crl, fimH, ial, papC* and *cjrC* is shown in Figure 5, and Supplementary Table 1. Among the identified VGs, *uid*A was present in all of the APEC phylotypes (100%) while *crl, fim*H and *ial* were found in 80 to 100% of the isolates examined. The abundance of *crl, fim*H and *ial* was 100.0% in the isolates of phylotype D2 of APEC while the prevalence of *pap*C and *cjr*C genes among these isolates was 77.78% and 77.22%, respectively (Figure 5). On the other hand, the prevalence of *pap*C gene were 50.0, 39.13 and 26.68% respectively in B2_2_, B2_3_ and A1 phylotypes of APEC, and *cjr*C gene was found in 41.67, 41.30 and 13.16% isolates of phylotypes B2_2_, B2_3_ and A1, respectively (Figure 5). However, none of the isolates of phylotype B1 possess these two (*pap*C and *cjr*C) genes. Thus, our present results revealed significant (p<0.05) association between two VGs and phylotypes (D2, B2). Conversely, only two phylotypes from healthy birds harbored only two VGs such as *papC* (phylotype A1) and *cjrC* (phylotype B2_3_) gene (Supplementary Table 2).

**FIGURE 5.**
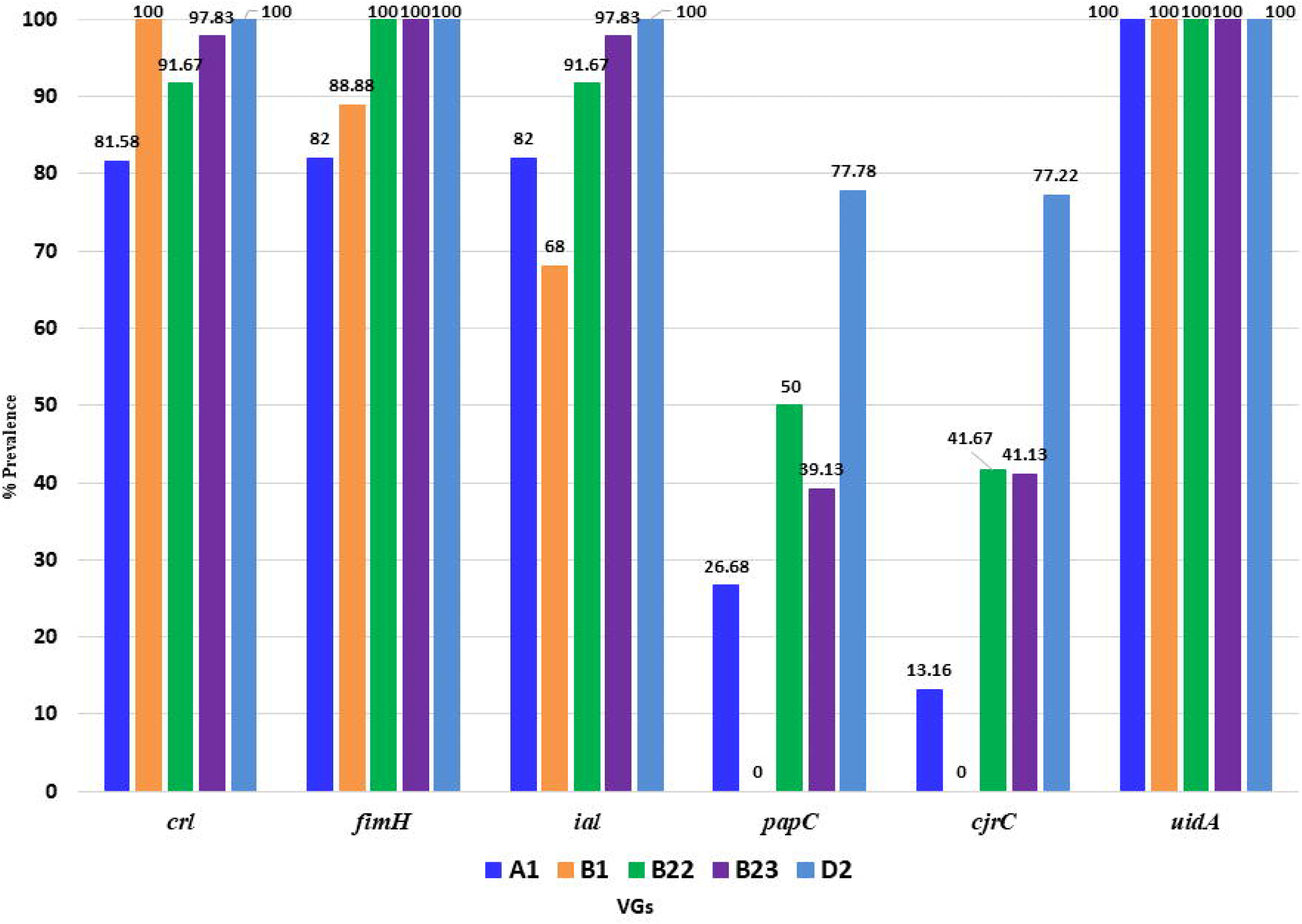
Prevalence of five virulent genes (VGs) among five APEC phylotypes. Here X axis represents the VGs while Y axis represents the prevalence (%) of the genes among the APEC isolates. For VG *crl*: the 1^st^ colored bar represents the prevalence of this gene in phylotype A1, 2^nd^ bar represents in phylotype B1, 3^rd^ bar represents in phylotype B2_2_, 4^th^ bar represents in phylotype B2_3_ and 5^th^ bar represents in phylotype D2. This serial is true for all the others VGs.

### 3.6 Genetic diversity of APEC isolates based on ribosomal (16S rRNA) gene sequencing

Nucleotide sequences obtained from 11 APEC isolates according to phylotyping and molecular typing (RAPD, ERIC and BOX-PCR) along with 13 previously reported reference sequences retrieved from NCBI database were used to generate a phylogenetic tree. The results obtained from the tree exhibited two different clades based on RAPD grouping and we referred to them as Clade A and Clade B which contained 5 and 6 isolates of *E. coli*, respectively (Figure 6). The strains of *E. coli* found in Clade A had 98-100.0% similarity with *E. coli* APEC O78 and *E. coli* EHEC strains; and strains found in clade B had closest relationship with *E. coli* O169:H41 strain. Interestingly, identified strains of *E. coli* from B2 and D2 phylotypes were found in both clades and the strains from phylogroups A1 and B1 were grouped into Clade A only.

**FIGURE 6.**
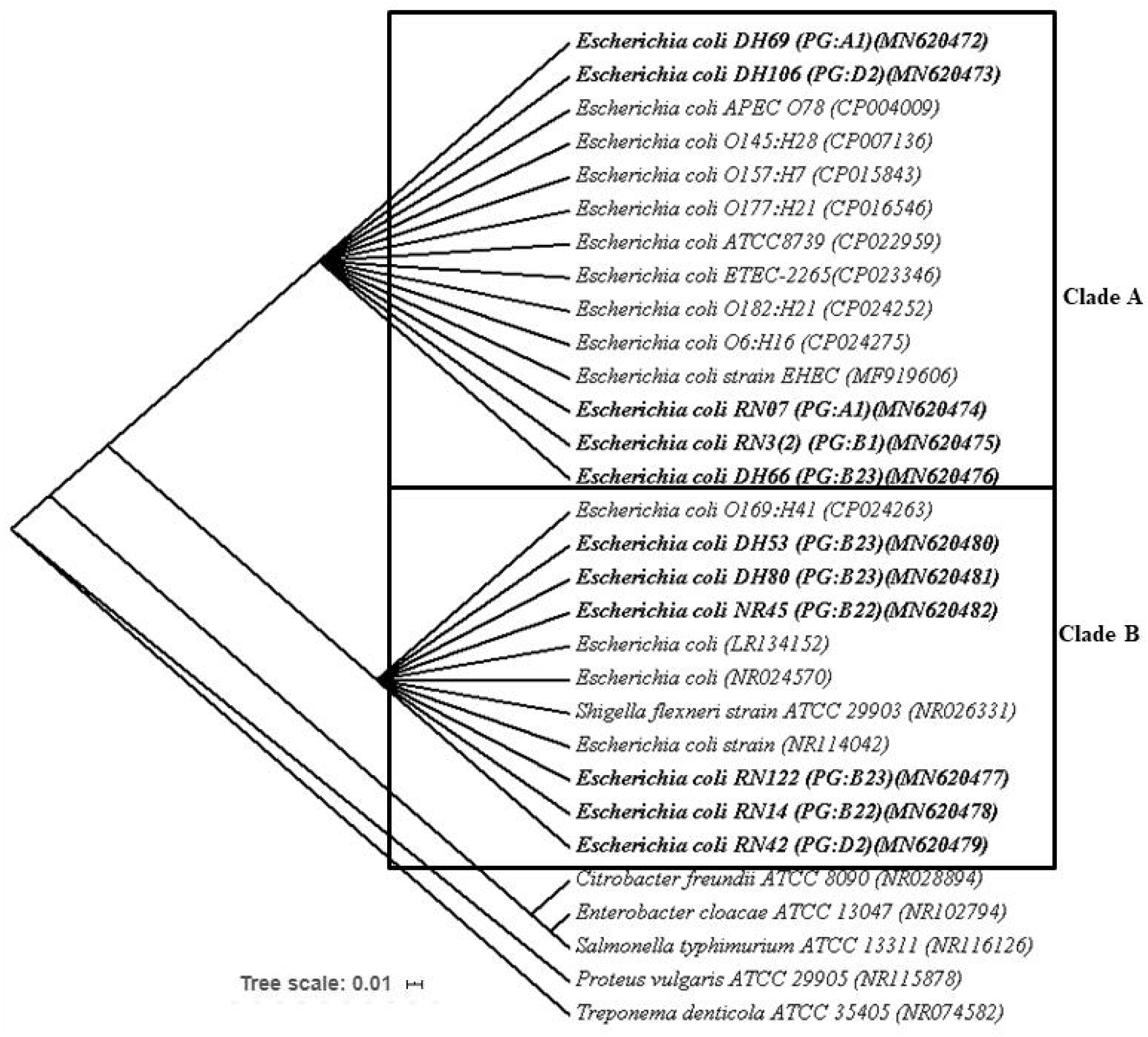
Phylogenetic tree predicted by the neighbor-joining method using 16S rRNA gene sequences. Kimura 2-parameter model method was used to compute the evolutionary distances., and The bootstrap considered 1000 replicates. The scale bar represents the expected number of substitutions averaged over all the analyzed sites. The optimal tree with the sum of branch length = 0.35475560 is shown here. *Treponema denticola* was used as out group. The length of the scale bar represents 1 nucleotide substitution per 100 positions.

### 3.7 Antibiogram profiling of circulating *E. coli* phylotypes

Antibiotic susceptibilities of randomly selected 114 isolates of *E. coli* were carried out using13 different antibiotics belonged to 11 groups (Table 3). The tested isolates showed a varying degree of resistance towards these antibiotics. The antibiogram profiling of the current study revealed that most *E. coli* isolates (78.95%) were resistant to doxycycline followed by nalidixic acid (76.32%), tetracycline (75.44%), ampicillin (74.76%), nitrofurantoin (63.16%), chloramphenicol (51.75%), cefoxitin (41.23%), ciprofloxacin (44.75%), sulfonamide (44.74%), azithromycin (31.58%), gentamycin (26.32%), imipenem (22.81%) and polymyxin B (7.78%). In regards to phylotypings, 61.96% isolates in phylotype B1 were resistant to at least three tested antibiotics, while, 55.24, 53.85, 51.16 and 45.58 % of the isolates of phylotypes D2, B2_2_, B2_3_ and A1, respectively were found to be resistant against 3≥ tested antibacterial agents (Table 1). However, in the comparative analysis, we found that 85.0% isolates belonging to phylotype A1 were resistant to tetracycline, and 77.5, 70.0 and 60.0% isolates of this phylotype were resistant to doxycycline, ampicillin and nalidixic acid, respectively. Resistance to tetracycline, doxycycline, nalidixic acid and ampicillin was 83.72, 79.07, 76.74 and 74.42% in isolates from phylotyping group B2_3_ and 81.82, 72.72, 72.72 and 90.91% in isolates from phylotype B2_2_, respectively (Table 1). The resistance tendency of the phylotype B1 to ampicillin, doxycycline, ciprofloxacin and gentamicin was higher than the resistance rate of the phylotype D2 but lower to tetracycline, nitrofurantoin, nalidixic acid and polymyxin B. Thus, on an average the isolates of B1, B2_2_ and D2 phylotypes were more resistant than those of phylotypes B2_3_ and A1 (Table 1). However, we did not find any significant differences (p>0.05) in MDR properties among the isolates of healthy and infected birds which indicated that indiscriminate and unauthorized use of antibiotics in the poultry farms of the Bangladesh. Furthermore, no significant differences were observed in the prevalence of VGs between nitrofurantoin-resistant and susceptible APEC strains except *papC*, remained more prevalent in the nitrofurantoin-susceptible strains (38.24%) than the nitrofurantoin-resistant strains (24%) (Supplementary Table 2).

**Table 1:**
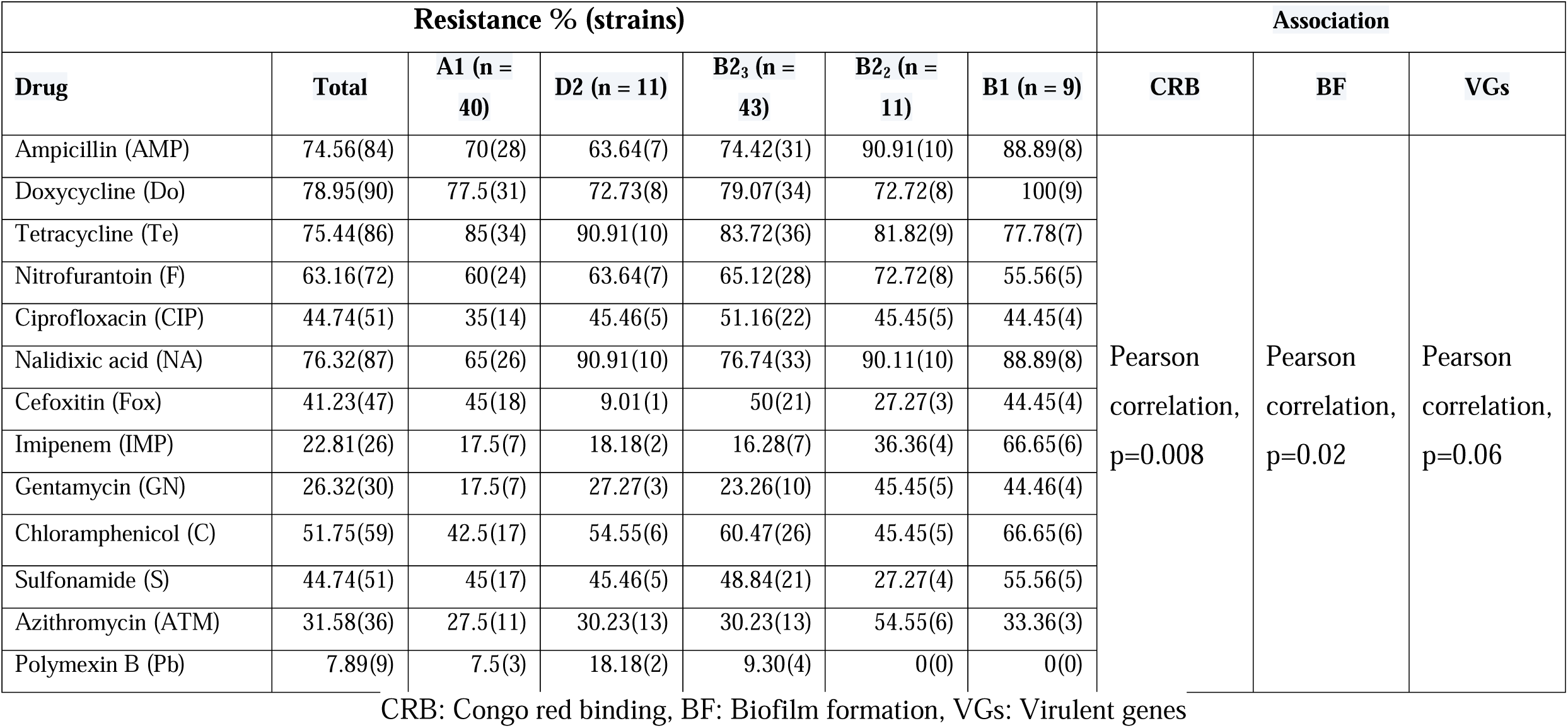
Relationship between *Escherichia coli* phylogenetic group and Drug sensitivity. Relationship between APEC Phylotypes and drug sensitivity.

| Resistance % (strains) |  |  |  |  |  |  | Association |  |  |
| --- | --- | --- | --- | --- | --- | --- | --- | --- | --- |
| Drug | Total | A1 (n = 40) | D2 (n = 11) | B2<sub>3</sub> (n = 43) | B2<sub>2</sub> (n = 11) | B1 (n = 9) | CRB | BF | VGs |
| Ampicillin (AMP) | 74.56(84) | 70(28) | 63.64(7) | 74.42(31) | 90.91(10) | 88.89(8) | Pearson correlation, p=0.008 | Pearson correlation, p=0.02 | Pearson correlation, p=0.06 |
| Doxycycline (Do) | 78.95(90) | 77.5(31) | 72.73(8) | 79.07(34) | 72.72(8) | 100(9) |  |  |  |
| Tetracycline (Te) | 75.44(86) | 85(34) | 90.91(10) | 83.72(36) | 81.82(9) | 77.78(7) |  |  |  |
| Nitrofurantoin (F) | 63.16(72) | 60(24) | 63.64(7) | 65.12(28) | 72.72(8) | 55.56(5) |  |  |  |
| Ciprofloxacin (CIP) | 44.74(51) | 35(14) | 45.46(5) | 51.16(22) | 45.45(5) | 44.45(4) |  |  |  |
| Nalidixic acid (NA) | 76.32(87) | 65(26) | 90.91(10) | 76.74(33) | 90.11(10) | 88.89(8) |  |  |  |
| Cefoxitin (Fox) | 41.23(47) | 45(18) | 9.01(1) | 50(21) | 27.27(3) | 44.45(4) |  |  |  |
| Imipenem (IMP) | 22.81(26) | 17.5(7) | 18.18(2) | 16.28(7) | 36.36(4) | 66.65(6) |  |  |  |
| Gentamycin (GN) | 26.32(30) | 17.5(7) | 27.27(3) | 23.26(10) | 45.45(5) | 44.46(4) |  |  |  |
| Chloramphenicol (C) | 51.75(59) | 42.5(17) | 54.55(6) | 60.47(26) | 45.45(5) | 66.65(6) |  |  |  |
| Sulfonamide (S) | 44.74(51) | 45(17) | 45.46(5) | 48.84(21) | 27.27(4) | 55.56(5) |  |  |  |
| Azithromycin (ATM) | 31.58(36) | 27.5(11) | 30.23(13) | 30.23(13) | 54.55(6) | 33.36(3) |  |  |  |
| Polymexin B (Pb) | 7.89(9) | 7.5(3) | 18.18(2) | 9.30(4) | 0(0) | 0(0) |  |  |  |

CRB: Congo red binding, BF: Biofilm formation, VGs: Virulent genes

### 3.8 Existence of Plasmid

Plasmid profiling of 45 randomly selected *E. coli* isolates based on their genetic diversity and antimicrobial resistance properties (Supplementary Table 2) showed that 73.33% (33/45) of *E. coli* isolates were plasmid bearing, and of them, 9.09% and 90.91% isolates belonged to healthy and diseased chicken’s samples, respectively (Figure 7, Supplementary Table 2). The molecular weight of plasmids varied from >30□kb to 2.1□kb. All of the plasmid harboring isolates showed multiple plasmid bands with size of 3 kb to 7.3 kb. However, the common size of plasmids was 3.9 kb to 5.6 kb as detected in all of the plasmid bearing strains (Figure 7). However, we demonstrated weak correlation (p=0.07) between plasmid bearing *E. coli* isolates with their phylogroups. With co-existence of the plasmid bearing genes and phylotypes, our results showed that 100 % isolates belonging to phylotype D2 of *E. coli* harbored plasmids of variable bands size and weight. On the other hand, 87.5, 83.33, 60 and 50% plasmid possessing isolates belonged to phylotypes B2_2_, B2_3_, B1 and A1 of *E. coli*, respectively (Figure 7, Supplementary Table 2).

**FIGURE 7.**
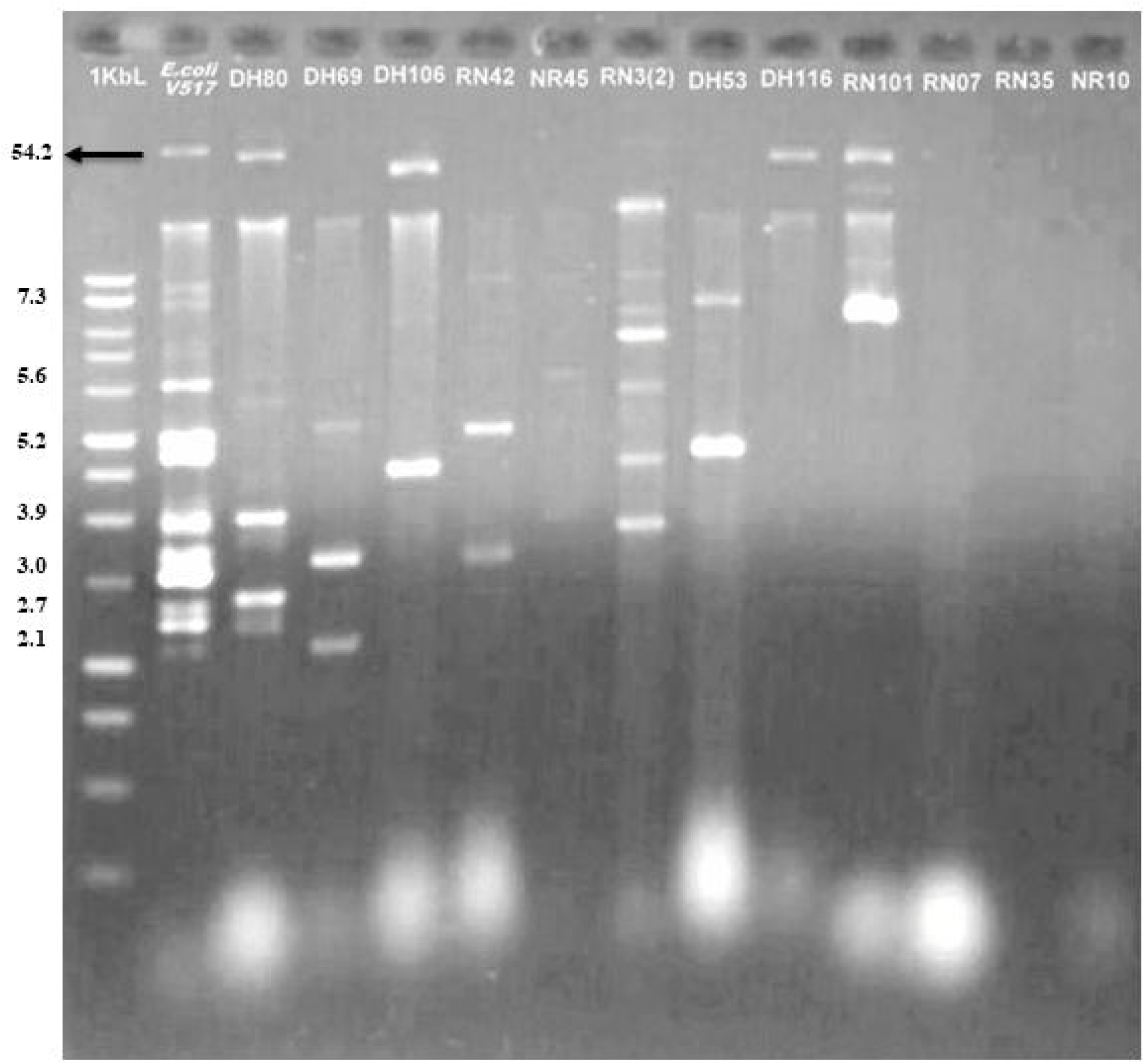
Plasmid profile of representative isolates of the multidrug resistant (MDR) *E. coli*: Agarose gel electrophoretogram (0.8% gel) of plasmid DNA of the MDR isolates: *E. coli* strain DH80, DH69, DH106, RN42, NR45, RN 3(2), DH53, DH116, RN07, RN35, NR10 (from lane 3 to lane 14), *E. coli* V517 in lane 2 was used as the marker and its plasmids of different sizes (kb) have been denoted in the figure. Lane 1 is molecular ladders. Plasmid bands at the same position was observed in *E. coli* strain DH80, RN3(2) at the position between 3.9 kb; strain DH69, RN42 at the positions 3.0 kb, strain DH53, RN101 at the position of about 7.3 kb and strain DH106, DH53, RN3(2) at the positions between 5.2 kb to 3.0 kb. Large plasmids at common positions were observed in strain DH80, DH106, DH116, RN101. Strain RN07, RN35, NR10 din not harbored any plasmid.

### 3.9 Associations between sample types, phylotyping and pathogenic intensity

The relationship analysis of the strength of sample types, molecular typing and pathogenic intensity was performed to determine possible associations among the isolates. In terms of molecular typing methods, we demonstrated that droppings (p=0.02), cloacal swabs (p=0.037) and internal organ liver (p=0.041) had significantly stronger correlation with RAPD followed by BOX (p=0.036, p=0.043, p=0.047, respectively), phylotypes (p=0.045, p=0.048, p=0.033, respectively) and ERIC (p=0.13, p=0.017, p=0.29, respectively) (Figure 8). While analyzing the pathogenicity and antibiogram profile of the tested isolates, sample categories had significantly higher correlation with CRB (p=0.008) followed by biofilm formation (p=0.02), drug sensitivity (p=0.03) and virulence genes (p=0.06). We also used correspondence analysis (CA) to measure the degree of relationship between the categories of the phylotypes, origin of samples (poultry feed and water, handler, egg surface, droppings, cloacal swab, liver) and pathogenic intensity (drug sensitivity). The bi-dimensional representation of phylotypes distribution in each of the seven sample categories is shown in Figure 9. The bi-dimensional representation explains 100% of the total variation with 68.55% explained by first dimension and 31.45% by the second dimension. On the other hand, the CA for the drug sensitivity and biofilm formation representation explains 87.25% variation by the first dimension and 12.75% by the second dimension. Moreover, in the current study correspondence analyses also demonstrated significant correlation among all of the phylotypes with Congo red binding (p=0.008), antimicrobial resistance (p=0.03), biofilm-formation (p=0.02), and virulent genes (p=0.06).

**FIGURE 8.**
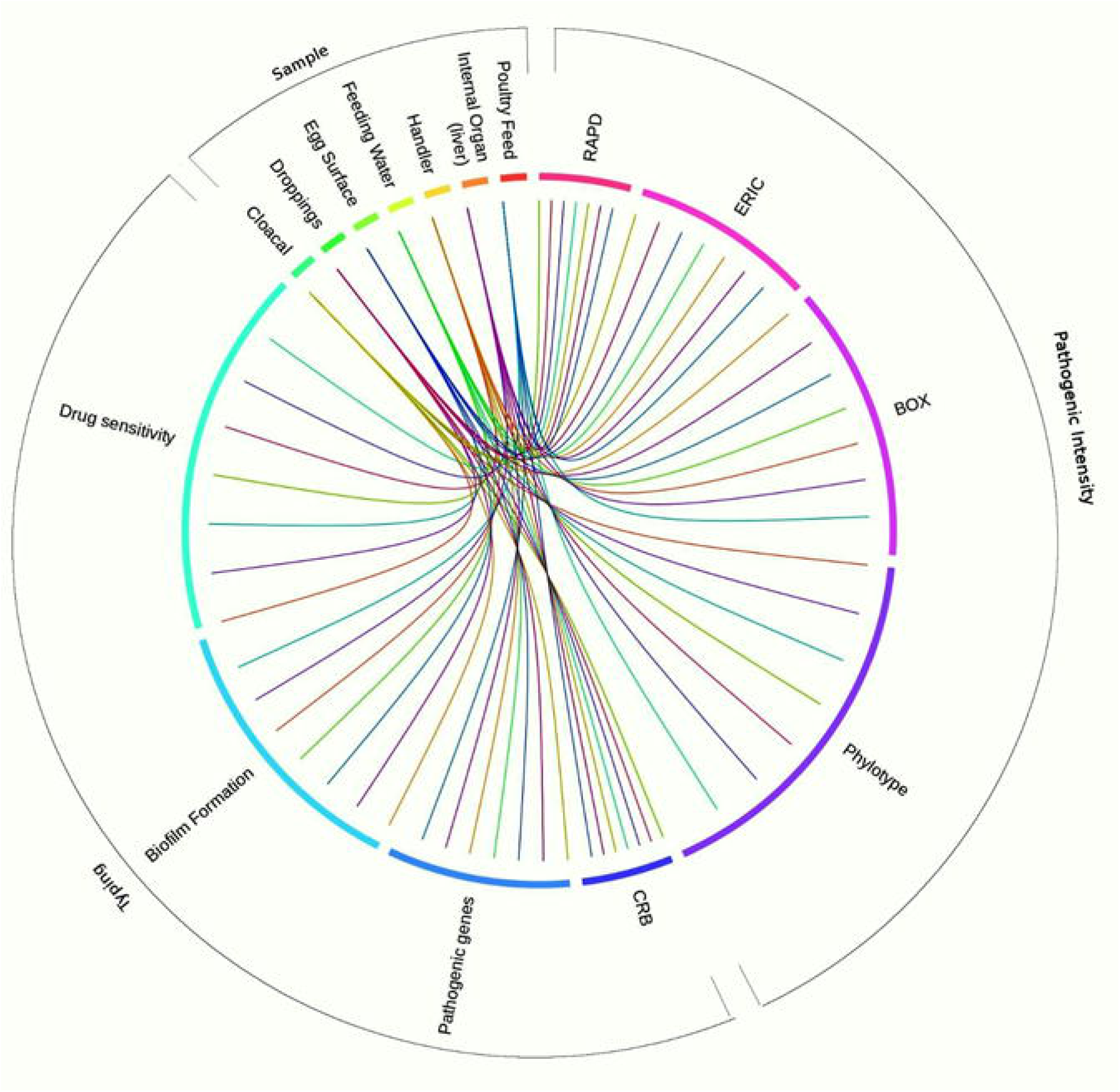
Circosplot representation of association of different sample types with molecular typing and pathogenic intensity characterization of *E. coli*. The association showed significant (*p*=0.032) correlation. The frequency of occurrence of different features such as sample categories, typing and pathogenicity patterns is depicted in the outer ring. The inner ring of Circos plot depicts the correlation between the sample categories (cloacal, droppings, egg surface, feeding water, handler, internal organ, feed), and molecular typing (RAPD, ERIC, BOX, phylotypes and pathogenic intensities (CRB, pathogenic genes, biofilm formation, drug sensitivity). The particular color is assigned to a particular color. The arc originates from sample types and terminates at typing, and pathogenic intensity levels to compare the association between the origin and terminating factors. The area of each colored ribbon depicts the frequency of the samples related with the particular typing and pathogenic intensity expression.

**FIGURE 9.**
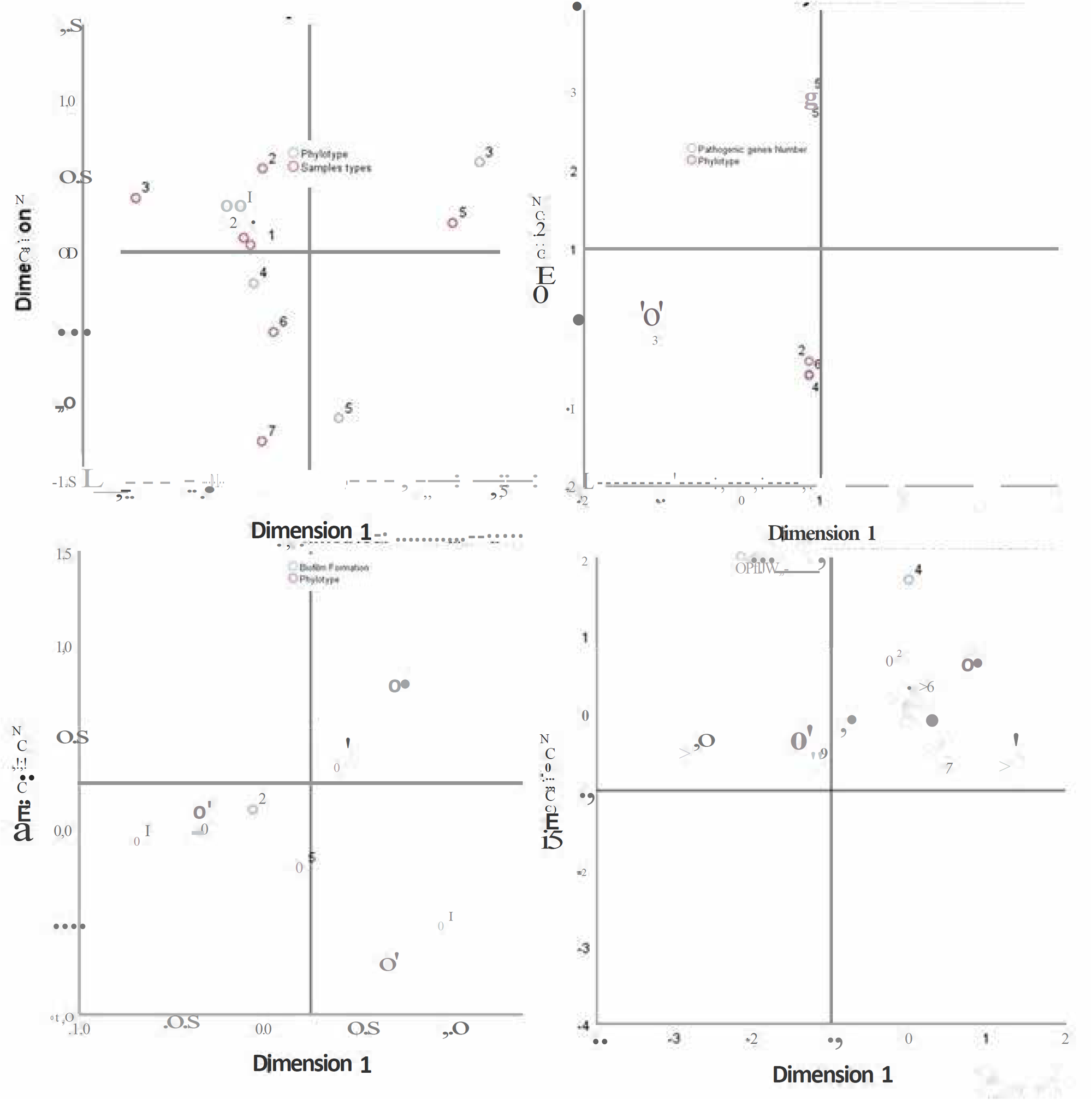
Correspondance analyses (CA) for the catégorial variables. Semple types, pathogenicity and phylotype that are similar where this two-dimensional representation explain 100% of the total variation, with 68.55% explained by 1^st^ dimension and 31.45% by the 2nd dimension. On the other hand, for the drug sensitivity and biofilm formation representation explains 87.25% variation by the 1st dimension and 12.75% by the 2^nd^ dimension. (a) Represents phylotypes (white circle) Vs sample types relation (red circle); (b) Represents phylotypes (red circle) Vs pathogenic gens relation (white circle); (c) Represents phylotypes (red circle) Vs biofilm formation (white circle); (a) Represents phylotypes (red circle) Vs drug sensitivity (white circle); Phylotypes: A1(1), B2_2_(2), B2_3_(3), B1(4), D2(5) (Supplementary table 2).

## 4. Discussion

The avian colibacillosis, caused by APEC is considered as one of the major threats to poultry industry and public health worldwide (Ibrahim et al., 2019). The devastating impacts of colibacillosis are particularly evident in the poultry farms of developing countries because of the poor hygiene practice, and management (Ebrahimi-Nik et al., 2018). In this study, we tested avian pathogenic *E. coli* (APEC) isolates from different poultry samples with regard to their phylotypes, phenotypic and genotypic virulence traits, and antimicrobial resistance. The findings of the current study provided evidence that the poultry farms could indeed be contaminated with multidrug-resistant (MDR) APEC phylotypes especially with the potentially pathogenic B2 and D2 phylotypes. This is particularly alarming for Bangladesh having a high disease burden, emergence of resistance traits, and the confluence of prevailing socio-economic, demographic and environmental factors (Azad et al., 2019; Sarker et al., 2019; Azad et al., 2017). This is the first study to report the association of multidrug-resistant APEC phylotypes in avian colibacillosis in different poultry farms of Bangladesh. A likely explanation to this high level of MDR in potentially pathogenic APEC isolates could be the improper disinfection management, lack of empty period of implementation between flocks, lack of knowledge about cleanliness, impure poultry feed and feeding environment, use of contaminated water and extensive use of antimicrobials in chickens, often without veterinary prescription, as reported in many earlier studies (Azad et al., 2019; Sarker et al., 2019; Azad et al., 2017; Reza et al., 2009). In the present study, 392 APEC isolates were obtained from 130 poultry samples (droppings, cloacal, feed, handler, egg, water, liver) collected from three districts (Narsingdi, Narayangonj and Manikgonj) of Bangladesh, with a clinical manifestation of colibacillosis at a prevalence rate of 44.39%. In Bangladesh, several earlier investigations in broiler chickens with colibacillosis reported 20 to 80% prevalence of this infectious disease (Rahman et al., 2017; Islam et al., 2014). The prevalence of colibacillosis in other countries like Nepal, China, Brazil and India ranged from 30 to 80% (Sarba et al., 2019; Saud et al., 2019; Younis et al., 2017; Chakraborty et al., 2015). The predisposing epidemiologic factors such as geographic locations, farm housing types, varying sample collection, transportation and preservation methods, and management practices are likely to contribute to the differences in frequency of pathogenic APEC isolation (Saud et al., 2019).

### 4.1 Circulatory molecular phylotypes of the APEC in Bangladesh

Phylogenetic analysis revealed that most of the isolates from phylotype B2_3_ followed by A1, D2, B2_2_ and B1 which provided a credible reference on the ecological distribution and genetic evolution of different pathogenic strains of APEC in the poultry farms of Bangladesh. Considering the elevated rate of APEC B2 and A1 phylotypes detected in this study and in consistent with previous report (Amer et al., 2018; Logue et al., 2017; Iranpour et al., 2015), we may infer that poultry samples could be a potential reservoir of APEC. Another study in Sri Lanka on phylogenetic diversity of APEC isolates from septicemic broiler and layer cases reported that the APEC isolates were belonged to A (71.00%), B1 (4.10%), B2 (7.90%) and D (18.70%) phylogroups (Dissanayake et al., 2008). However, most of the recent studies reported phylotypes A and D as the most abundant phylotypes of AEPC isolated from poultry such as in Italy (Pasquali et al., 2015), China (Wang et al., 2010), Canada (Aslam et al., 2014), and Iran (Ghanbarpour et al., 2011) indicating that the frequency of phylotypes might vary among different geographic regions. Globally, phylotypes B2 and D are classified as pathogenic *E. coli* (Clermont, et al., 2013), and our present findings are in accordance to these findings. Although recent studies that utilize the updated method by Clermont et al. (2013) are scarce, Logue et al. (2017) classified APEC isolates according to the new phylogenetic typing and concluded that strains in A and B1 group were of lower pathogenic potential. The low prevalence of phylogenetic group D2 was also reported earlier in poultry (Pasquali et al., 2015; Ghanbarpour et al, 2011). In addition to phylotyping, three molecular typing methods such as RAPD, ERIC and BOX PCR were used to reveal the genetic relatedness among the APEC isolates (Daga et al., 2003). Though MLST technique is one of the important and widely used tool for APEC characterization globally, we did not utilize this technique in APEC phylotyping considering some of its inherent disadvantages (Johnson et al., 2017; Salipante et al., 2015).

### 4.2 Pathogenic properties of the circulating APEC

APEC isolates often carried a broad range of virulence genes (VGs) that may enable their pathogenicity in avian colibacillosis. These include production of adhesions, toxins, siderophores, iron transport systems, and invasins (Amer et al., 2018; Borzi et al., 2018; Ahmed and Shimamoto, 2013, Dziva and Stevens, 2008). Several VGs such as *ial, pap*C, *fim*H, *crl* are important in APEC adherence (Borzi et al., 2018). In this study, many of the APEC isolates belonged to A1, B1, B2 and D2 phylotypes carried one or more virulence genes which were represented by *pap*C, *cjr*C, *crl, ial, fim*H and *uid*A. The phylogroup B2 and D2 (pathogenic strains) harbored all of these virulence determinants while phylotype A and B1 (usually found in commensal strains) possessed few VGs. Smith et al. (2007) reported that the phylotype B2 and D possessed several pathogenicity associated islands, and express multiple virulence factors such as adherence factors including biofilm production supporting our current investigation. In the present investigation, we indicated that the distribution of 6 VGs was different, and a large proportion of adherence genes had strong biofilm formation ability. This also revealed a positive correlation among these genes, biofilm phenotype and phylotypes. In the current study, the VGs *uid*A, *crl, fimH and ial* were found in 80 to 100% of the APEC isolates examined. These results are in accordance with many of the previously published studies (Amer et al., 2018; Reichhardt et al., 2018; Reichhardt et al., 2015; Ahmed and Shimamoto, 2013; Johnson et al., 2008). Moreover, the presence of similar VGs in APEC isolates proposed that APEC isolates can act as zoonotic pathogens and reservoirs of virulence causing human infections (Mora et al., 2013). The *fim*H virulence factor is seemed to be an essential unit for protecting the APEC isolates against host immune system but the exact role of *fim*H in the pathogenicity of APEC isolates remains debatable with incompatible results (Asadi et al., 2018). Congo red binding (CRB) assay has been used extensively to supplement nutrient agar as a selection medium to distinguish curli-producing bacteria from non-curliated bacteria when CR-binding is confirmed to be curli-dependent (Reichhardt et al., 2015; Amer et al., 2018). Furthermore, curli fibers are protease resistant and bind to Congo red (CR) and other amyloid dyes (Reichhardt and Cegelski, 2018). The curli-negative mutant isolates have less adherence colonization, invasion and persistence to chicken tissues recommending curli as a virulence factor (Reichhardt and Cegelski, 2018; Reichhardt et al., 2015; Mokady et al., 2005). Therefore, it can be supposed that most of APEC isolates are curliated (Reichhardt and Cegelski, 2018; Reichhardt et al., 2015; Mora et al., 2013). The relative abundance of *cjrC* gene seemed to be positively correlated with antimicrobial resistance profile of the isolates of phylotypes D2, B2_3_ and A1. In another study, Zhao et al. found that the prevalence of iroN gene (Salmochelins related) in Cefoxitin susceptible UPEC isolates was significantly higher than those resistant ones, and nitrofurantoin-resistant isolates had reduced VGs compared with susceptible strains (Zhao et al., 2009). The *pap*C genes however had relatively lower abundance among the APEC isolates, and we did not reveal any differences in the prevalence of VGs between nitrofurantoin-resistant and susceptible APEC strains. This result is line with the findings of many other studies (Sgariglia et al., 2019; Borzi et al., 2018; Zhao et al., 2009). Furthermore, most of the virulence determinants identified in this study could be acquired by horizontal transmission without disrupting the clonal lineage.

Therefore, the most likely reason for the diversity in distribution of these virulence determinants in different phylogenetic groups is horizontal gene acquisition. Nevertheless, it is yet unknown whether the mere acquisition of these genes is enough to make an organism virulent or if a specific genetic background is required for the transfer and expression of these genes (Dissanayake et al., 2008). Moreover, biofilm formation is an important virulence factor for APEC strains and contributes to the resistance to different classes of antimicrobials (Hoque et al., 2020). APEC strains identified in this study showed broad spectrum of antimicrobial resistance, and possessed biofilm forming abilities, which might be the potential factors for Colibacillosis in the poultry farm, and persistence of the disease, and increased risk of transmission to non-infected birds. However, pathogenic potentials of the APEC associated VGs have not been demonstrated using *in vivo* animal trials which is one of the drawbacks of the current study.

### 4.4 Correlations between phylotypes and MDR of circulating APEC

Colibacillosis in the poultry farms might be prevented and/or controlled by the rational therapeutic use of antimicrobials. However, evolution of MDR APEC strains along with the transmission of resistance genes has created challenges in reducing risk of APEC infections (Subedi et al., 2018). Regarding antimicrobial resistance exhibited by the isolates of different phylotypes, our results indicated that phylotype A1 isolates were more susceptible than the isolates of other phylotypes. Conversely, the isolates of phylotype B1 displayed the highest antimicrobial resistance pattern. Varying results have been reported by other investigators, indicating that although being more virulent, the isolates of phylotype B2 were more susceptible to antibiotics (Chakraborty et al., 2015). However, we found that resistance to doxycycline, ampicillin, and nalidixic acid was common in group B2 isolates while the phylotype A1 remained resistant to tetracycline only. Therefore, our present findings demonstrated that strains belonged to phylotypes B1, D2 and B2_2_ were carrying more resistance and/or virulent properties than the strains of phylotype B2_3_ and A1 corroborating the findings of Iranpour et al. (2015) and Moreno et al. (2006). In this study, we demonstrated that all of the APEC phylotypes possessed MDR properties which did not comply with the previous findings of Etebarzadeh et al. and Iranpour et al. who reported that only the phylotype B2 of APEC isolates could bear MDR phenomena (Iranpour et al.,2015; Etebarzadeh et al., 2012). This variation could be explained by the horizontal transfer of resistance genes through plasmids across the APEC strains. These findings therefore imply that in the poultry industry of Bangladesh, a large number of poultry samples might act as a reservoir for such resistant strains. Unfortunately, we did not find any single APEC isolate showing sensitivity to all of the 13 antibiotics tested, which might be due to the widespread, indiscriminate and long-term use of similar drugs in the poultry farms (Subedi et al., 2019; Li et al., 2015). None the less, all of the (100.0%) plasmid bearing strains found in this study were multi-drug resistant (MDR). Even though, isolates which did not bear any plasmid DNA (26.67%) were also resistant to at least three or more antimicrobials tested (Prescott et al., 2000), which could make it more possible for a susceptible bacterium to acquire resistance factors through conjugation or transformation (Miles et al., 2006). However, the number of plasmids found in a particular isolate would not necessarily indicate the level of MDR properties of the isolate which might also be one of the predisposing causes of spreading and developing MDR properties among poultry population in Bangladesh.

### 4.5 Correlations between sample types, molecular typing methods, pathogenic intensity and MDR properties of circulating APEC

The phylotype distribution of circulating APEC isolates was influenced by many factors such as sample types, CRB, biofilm formation (BF) and VGs (Ramadan et al., 2016). The correspondence analysis (CA) and circular visualization revealed stronger association between pathogenic intensity (CRB, BF, VGs) and molecular typing (phylotyping), and these phenomena of APEC isolates might be associated to MDR properties of this bacterium in the poultry farms of Bangladesh as supported by previous studies (Wang et al., 2016; Coura, et al., 2017; Carlos et al., 2010). Our results indicated that phylotype B2 was the main circulating APEC followed by phylotype A1 (Figures 8 & 9, Supplementary Table 2). However, both the results of CA and circular plot also showed that APEC phylotypes B2, B1, D2 and A1 were predominantly isolated from droppings and cloacal samples. Though, the CA analysis did not display stronger association between B2 and D2 APEC phylotypes, their BF ability and VGs, however, further visualization using the circular plot showed that these two phylotypes had higher potentials for BF, and harbored more VGs than other APEC phylotypes identified in this study (Figures 8 & 9, Supplementary Table 2). These findings also corroborates with many earlier studies (Wang et al., 2016; Carlos et al., 2010). Furthermore, both CA and circular plot visualization showed that all of the APEC phylotypes isolated from different poultry samples possessed MDR phenomena as also reported in many recent studies (Subedi et al., 2019; Li et al., 201;). Therefore, high prevalence of antibiotic resistant APEC strains, and their associations with sample types, molecular typing methods, pathogenic intensity and MDR properties suggest an alternative approach of organic antimicrobial compounds, and/or metals usages (Hoque et al., 2020), and the rotational selection and judicious use of antibiotics in the poultry farms in Bangladesh.

## 5. Conclusions

The identification of virulence genes (VGs) and antimicrobials resistance from diverge samples of avian colibacillosis reveals the great importance of APEC zoonotic potential. Our results showed that five phylogroups were prevailing among the APEC isolates, and of them, phylotypes A1 and B2 were the most common groups. Phylogenetic analysis revealed two distinct clades (Clade A and Clade B) of APEC. Phylogroups B2 and D2 were found in strains both clades, and phylogroups A1 and B1 were found in clade A only. Antibiogram profiling revealed that 100.0% isolates, and majority of the phylotypes (> 50.0) B1, D2, B2_2_, B2_3_ and A1 were resistant to at least antibiotics. Our data demonstrated that relatively high MDR levels among all of the APEC phenotypes is a serious concern in Bangladesh, and may led to several adverse effects on animals, humans, and the environment. Furthermore, the APEC isolates and their associated phylogroups in the present study harbored a set of VGs in a relatively higher proportion along with potential BF ability. These two phenomena of the APEC phylotypes could be associated to their MDR properties, which could cause a serious public health problem. However, future research should be done to better understand the flow of antimicrobials usage, monitoring the spread of ARGs and VGs using larger population size in different ecosystems of the poultry sectors of Bangladesh.

## Supporting information

Main Figure Legends

## Author’s contributions

O.S. carried out the studies (sampling, sequencing, molecular and data analysis). O.S. and O.K.I. carried out the biofilm assays. M.N.H. performed the statistical and correspondence analyses. O.S. and M.N.H drafted the manuscript. M.M.R, M.S. and M.A.H. developed the hypothesis, supervised the whole work and critically review the drafted manuscript. All authors read and approved the final manuscript.

## Acknowledgments

The authors would like to acknowledge M. Al Amin, PhD fellow, Department of Microbiology, University of Dhaka for his assistance in sampling. We would like to acknowledge Bangabandhu Science & Technology Fellowship Trust for supporting Otun Saha with PhD fellowship.

## Data availability

The 16S rRNA gene sequencing data (11 sequence) has been submitted to NCBI database under the accession numbers-MN620472-MN620482.

## Ethical approval

The Animal Experimentation Ethical Review Committee (AEERC), Faculty of Biological Sciences, University of Dhaka ethically approve research work under Reference No. 71/Biol.Scs./2018-2019.

## Supplementary Figure Legends

**SUPPLEMENTARY FIGURE 1**| RAPD patterns of bacterial isolate using primer 1283. Lane 2 is negative blank control and lanes1 is molecular ladders. Lanes 3–18 are samples DH80, DH69, DH66, RN07, DH106, RN14, RN3 (2), RN50, RN42, DH116, NR10, NR45, DH53, RN101, NR34, RN122 respectively representing group 1-10.

**SUPPLEMENTARY FIGURE 2**| ERIC-PCR patterns of bacterial isolate using primer ERIC1 and ERIC2. Lane 2 is negative blank control and lanes1 is molecular ladders. Lanes 3–17 are samples DH80, DH69, DH66, RN122, DH53, DH66, RN07, RN93, DH106, RN14, RN3 (2), RN101, RN42, NR45, RN88 respectively representing group 1-8.

**SUPPLEMENTARY FIGURE 3**| BOX-PCR patterns of bacterial isolate using primer BOXA1R. Lane 2 is negative blank control and lanes 1 is molecular ladders. Lanes 3–17 are samples DH80, RN14, DH69, DH66, DH3, NR35, RN07, DH106, RN3 (2), RN42, NR45, RN93, NR48, RN122, RN89 respectively representing group 1-9.

**SUPPLEMENTARY FIGURE 4**| PCR results for the detection of *E. coli* virulent genes (VGs) among the colibacillosis cases of Bangladeshi poultry samples. A) (*uid*A: 147bp) Lane 2 is negative blank control and lanes 1 is molecular ladders (1kb). Lanes 3-11 are strain DH53, DH69, DH66, DH80, DH106, NR45, RN122, RN07, RN88. B) (*crl*: 250bp) Lane 2 is negative blank control and lanes 1 is molecular ladders (100bp). Lanes 3–10 are strain DH53, DH69, DH80, NR45, RN07, RN3 (2). C) (*pap*C: 328bp) Lane 2 is negative blank control and lanes 1 is molecular ladders (100bp). Lanes 3–4 are strain DH69, RN101. D) (*ial*: 650bp) Lane 2 is negative blank control and lanes 1 is molecular ladders (1kb). Lanes 3-10 are strain DH53, DH69, DH80, DH106, NR45, RN101, RN07, DH66.E) (*fim*H: 164bp) Lane 2 is negative blank control and lanes 1 is molecular ladders (1kb). Lanes 3-12 are strain DH53, DH69, DH66, DH106, DH80, RN122, RN42, RN07, RN3 (2), NR45.F) (*cjr*C: 518bp) Lane 6 is negative blank control and lanes 5 is molecular ladders (100bp). Lanes 1-4 are strain DH106, RN83, NR45, NR6.

## Supplementary Tables

**Supplementary Table 1:** Sequence of oligonucleotide primers of different target genes used in this study to detect pathogenic *Escherichia coli* strains.

**Supplementary Table 2:** Relative comparison among isolated Pathogenic *Escherichia coli* from different poultry farms of three sampling locations

## References

Abbey, T. C., & Deak, E. (2019). What’s New from the CLSI Subcommittee on Antimicrobial Susceptibility Testing M100. Clin. Microbiol. News. 41(23), 203–209

Ahmad, M.D. et al. (2009). Drinking water quality by the use of Congo Red medium to differentiate between pathogenic and nonpathogenic E. coli at poult. farms. J. Anim. Plant Sci. 19, 108–110.

Ahmed, A. M., Shimamoto, T., & Shimamoto, T. (2013). Molecular characterization of multidrug-resistant avian pathogenic *Escherichia coli* isolated from septicemic broilers. Int. J. Med. Microbiol. 303(8), 475–483.

Al Azad, M., Rahman, A., Rahman, M., Amin, R., Begum, M., Ara, I. et al. (2019). Susceptibility and Multidrug Resistance Patterns of *Escherichia coli* Isolated from Cloacal Swabs of Live Broiler Chickens in Bangladesh. Pathogens 8(3), 118.

Amer, M. M., Bastamy, M. A., Ibrahim, H. M., Salim, M. M. (2015). Isolation and characterization of avian pathogenic Escherichia coli from broiler chickens in some Governorates of Egypt. Vet. Med. J.–Giza, 61–1.

Amer, M. M., Mekkym H. M., Amer, A. M., Fedawy, H.S. (2018) Antimicrobial resistance genes in pathogenic Escherichia coli isolated from diseased broiler chickens in Egypt and their relationship with the phenotypic resistance characteristics. Vet. World 11(8), 1082–1088.

Asadi, A., ZahraeiSalehi, T., Jamshidian, M., & Ghanbarpour, R. (2018). ECOR phylotyping and determination of virulence genes in *Escherichia coli* isolates from pathological conditions of broiler chickens in poultry slaughter-houses of southeast of Iran. Vet. Res. Forum: an int. quarterly J. 9(3), 211–216.

Aslam, M., Toufeer, M., Bravo, C. N., Lai, V., Rempel, H., Manges, A., & Diarra, M. S. (2014). Characterization of extraintestinal pathogenic *Escherichia coli* isolated from retail poultry meats from Alberta, Canada. Int. J. Food Microbiol. 177, 49–56.

Azad, M.ara, Amin, R., Begum, M. I. A., Fries, R., Lampang, K. N., & Hafez, H. M. (2017). Prevalence of antimicrobial resistance of *Escherichia coli* isolated from broiler at Rajshahi region, Bangladesh. Br. J. Biomed. Multidiscip. Res. 1(1), 6–12.

Bikandi, J., Millán, R. S., Rementeria, A., & Garaizar, J. (2004). In silico analysis of complete bacterial genomes: PCR, AFLP–PCR and endonuclease restriction. Bioinformatics 20(5), 798–799.

Bist, B., Sharma, B., & Jain, U. (2014). Virulence associated factors and antibiotic sensitivity pattern of *Escherichia coli* isolated from cattle and soil. Vet. World 7(5).

Borzi, M. M., Cardozo, M. V., de Oliveira, E. S., de Souza Pollo, A., Guastalli, E. A. L., dos Santos, L. F., & de Ávila, F. A. (2018). Characterization of avian pathogenic Escherichia coli isolated from free-range helmeted guineafowl. Brazilian J. Microbiol. 49, 107–112.

Carlos, C., Pires, M. M., Stoppe, Johnson N. C., Hachich, E. M., Sato, M. I., Gomes, T. A. & Ottoboni, L. M. (2010). *Escherichia coli* phylogenetic group determination and its application in the identification of the major animal source of fecal contamination. BMC Microbiol. 10(1), 161.

Chakraborty, A., Saralaya, V., Adhikari, P., Shenoy, S., Baliga, S., & Hegde, A. (2015). Characterization of *Escherichia coli* Phylogenetic Groups Associated with Extraintestinal Infections in South Indian Population. Annals Med. Health Sci. Res. 5(4), 241–246.

Clermont, O., Christenson, J. K., Denamur, E., & Gordon, D. M. (2013). The Clermont *Escherichia coli* phyloLJtyping method revisited: improvement of specificity and detection of new phyloLJgroups. Env. Microbiol. Reports. 5(1), 58–65.

Coura, F. M., Diniz, S. A., Silva, M. X., Arcebismo, T. L., Minharro, S., Feitosa, A. C., & Heinemann, M. B. (2017). Phylogenetic group of *Escherichia coli* isolates from broilers in Brazilian Poultry Slaughterhouse. The Sci. World J. 7.

Daga, A. P., Koga, V. L., Soncini, J. G. M., de Matos, C. M., Perugini, M. R. E., Pelisson, M. & Vespero, E. C. (2019). *Escherichia coli* Bloodstream Infections in Patients at a University Hospital: Virulence Factors and Clinical Characteristics. Front. Cell. Infect. Microbiol. 9(191).

De Campos, T. A., Stehling, E. G., Ferreira, A., de Castro, A. F. P., Brocchi, M., & da Silveira, W. D. (2005). Adhesion properties, fimbrial expression and PCR detection of adhesin-related genes of avian *Escherichia coli* strains. Vet. Microbiol. 106(3-4), 275–285.

De Carli, S., Ikuta, N., Lehmann, F. K. M., da Silveira, V. P., de MeloPredebon, G., Fonseca, A. S. K. & Lunge, V. R. (2015). Virulence gene content in *Escherichia coli* isolates from poultry flocks with clinical signs of colibacillosis in Brazil. Poult. Sci. 94(11), 2635–2640.

Dissanayake, D. R. A., Wijewardana, T. G., Gunawardena, G. A. & Poxton, I. R. (2008). Distribution of lipopolysaccharide core types among avian pathogenic *Escherichia coli* in relation to the major phylogenetic groups. Vet. Microbiol. 132(3-4), 355–363.

Dombek, P. E., Johnson, L. K., Zimmerley, S. T. & Sadowsky, M. J. (2000). Use of repetitive DNA sequences and the PCR to differentiate *Escherichia coli* isolates from human and animal sources. Appl. Environ. Microbial. 66(6), 2572–2577.

Dziva, F. & Stevens, M. P. (2008). Colibacillosis in poultry: unravelling the molecular basis of virulence of avian pathogenic *Escherichia coli* in their natural hosts. Avian Patho. 37(4), 355–366.

Ebrahimi-Nik H., Bassami, M.R., Mohri, M., Rad, M. & Khan, M.I. (2018) Bacterial ghost of avian pathogenic *E. coli* (APEC) serotype O78:K80 as a homologous vaccine against avian colibacillosis. PLOS One13 (3), e0194888.

Etebarzadeh, Z., Oshaghi, M. & Mozafari, N. A. (2012). Evaluation of relationship between phylogenetic typing and antibiotic resistance of uropathogenic *Escherichia coli*. J. Microbiol. World 3-4(11), 84–92.

Ezz El Deen, A. et al. (2010). Characterization of Surface Proteins of Escherichia coli Isolated from Different Egyptian Sources. Int. J. Microbiol. Res. 1, 147–161.

Feng, P., Weagant, S. D., Grant, M. A., & Burkhardt, W. (2015). Bacteriological Analytical Manual Chapter 4 Enumeration of *Escherichia coli* and the Coliform Bacteria, U.S. Food and Drug Administration 10903 New Hampshire Avenue Silver Spring, MD 20993 1-888-INFO-FDA (1-888-463-6332).

Ghanbarpour, R., Sami, M., Salehi, M. & Ouromiei, M. (2011). Phylogenetic background and virulence genes of *Escherichia coli* isolates from colisepticemic and healthy broiler chickens in Iran. Trop. Anim. Health Prod. 43(1), 153–157.

Glass-Kaastra, S. K., Pearl, D. L., Reid-Smith, R., McEwen, B., Slavic, D., Fairles, J. & McEwen, S. A. (2014). Multiple-class antimicrobial resistance surveillance in swine *Escherichia coli* F4, *Pasteurella multocida* and *Streptococcus suis* isolates from Ontario and the impact of the 2004–2006 Porcine Circovirus type-2 Associated Disease outbreak. Prev. Vet. Med. 113(2), 159–164.

Hojati, Z., Zamanzad, B., Hashemzadeh, M., Molaie, R. & Gholipour, A. (2015). The *fim* H gene in uropathogenic *Escherichia coli* strains isolated from patients with urinary tract infection. Jundishapur J. Microbiol. 8(2).

Hoque M. N. et al. (2018). Molecular characterization of *Staphylococcus aureus* strains in bovine mastitis milk in Bangladesh. Int. J. Vet. Sci. Med., 6(1), 53–60.

Hoque M. N. et al. (2019). Metagenomic deep sequencing reveals association of microbiome signature with functional biases in bovine mastitis. Sci. Rep. 9, 13536.

Hoque M. N. et al. (2020). Insights into the Resistome of Bovine Clinical Mastitis Microbiome, a Key Factor in Disease Complication. Front. Microbiol. 11, 860, doi: 10.3389/fmicb.2020.00860.

Houser, B. A., Donaldson, S. C., Padte, R., Sawant, A. A., DebRoy, C. & Jayarao, B. M. (2008). Assessment of phenotypic and genotypic diversity of *Escherichia coli* shed by healthy lactating dairy cattle. Foodborne Patho. Dis. 5(1), 41–51.

Hu, Y., Yan, C., Hsu, C. H., Chen, Q. R., Niu, K., Komatsoulis, G. A. & Meerzaman, D. (2014). OmicCircos: a simple-to-use R package for the circular visualization of multidimensional omics data. Cancer Inform. 13, CIN–S13495.

Ibrahim, R. A., Cryer, T. L., Lafi, S. Q., Basha, E. A., Good, L. & Tarazi, Y. H. (2019). Identification of *Escherichia coli* from broiler chickens in Jordan, their antimicrobial resistance, gene characterization and the associated risk factors. BMC Vet. Res. 15(1), 159.

Iranpour, D., Hassanpour, M., Ansari, H., Tajbakhsh, S., Khamisipour, G. & Najafi, A. (2015). Phylogenetic groups of *Escherichia coli* strains from patients with urinary tract infection in Iran based on the new Clermont phylotyping method. BioMed Re. Int. 846219, doi: 10.1155/2015/846219.

Islam, M. M., Islam, M. N., Sharifuzzaman, F. M., Rahman, M. A., Sharifuzzaman, J. U. et al. (2014). Isolation and identification of *Escherichia coli* and *Salmonella* from poultry litter and feed. Int. J. Nat. Soc. Sci. 1(1), 1–7.

Jang, J., Hur, H. G., Sadowsky, M. J., Byappanahalli, M. N., Yan, T. & Ishii, S. (2017). Environmental *Escherichia coli*: ecology and public health implications—a review. J. Applied Microbiol. 123(3), 570–581.

Johnson, J. R., Johnston, B. D. & Gordon, D. M. (2017). Rapid and Specific Detection of the *Escherichia coli* Sequence Type 648 Complex within Phylogroup F. J. Clinical Microbiol. 55 (4), 1116–1121.

Johnson, J. R., Kuskowski, M. A., Owens, K., Gajewski, A. & Winokur, P. L. (2003). Phylogenetic origin and virulence genotype in relation to resistance to fluoroquinolones and/or extended-spectrum cephalosporins and cephamycins among *Escherichia coli* isolates from animals and humans. The J. Infect. Dis. 188(5), 759–768.

Johnson, T. J., Wannemuehler, Y., Doetkott, C., Johnson, S. J., Rosenberger, S. C. & Nolan. L. K. (2008). Identification of minimal predictors of avian pathogenic *Escherichia coli* virulence for use as a rapid diagnostic tool. J. Clin. Microbiol. 46:3987–3996.

Kabiswa, W., Nanteza, A., Tumwine, G. & Majalija, S. (2018). Phylogenetic Groups and Antimicrobial Susceptibility Patterns of *Escherichia coli* from Healthy Chicken in Eastern and Central Uganda. J. Vet. Med. 9126467, doi: 10.1155/2018/9126467.

Knöbl, T. et al. (2011). Serogroups and virulence genes of Escherichia coli isolated from psittacine birds. Pesq. Vet. Bras. 31(10), 916–921.

Knöbl, T., Moreno, A. M., Paixão, R., Gomes, T. A., Vieira, M. A., da Silva Leite, D. & Ferreira, A. J. (2012). Prevalence of avian pathogenic *Escherichia coli* (APEC) clone harboring *sfa* gene in Brazil. The Sci. World J. 437342.

Kumar, S., Stecher, G. & Tamura, K. (2016). MEGA7: molecular evolutionary genetics analysis version 7.0 for bigger datasets. Mol. Biol. Evol. 33(7), 1870–1874.

Kur, J. W. (2009). Biofilm formation as a virulence determinant of uropathogenic *Escherichia coli* Dr+ strains. Polish Society Microbiol. 58(3), 223–229.

Li, Y., Chen, L., Wu, X. & Huo, S. (2015). Molecular characterization of multidrug-resistant avian pathogenic *Escherichia coli* isolated from septicemic broilers. Poult. Sci. 94(4), 601–611.

Logue, C. M., Wannemuehler, Y., Nicholson, B. A., Doetkott, C., Barbieri, N. L. & Nolan, L. K. (2017). Comparative analysis of phylogenetic assignment of human and avian ExPEC and fecal commensal *Escherichia coli* using the (previous and revised) Clermont phylogenetic typing methods and its impact on avian pathogenic *Escherichia coli* (APEC) classification. Front. Microbiol. 8, 283.

López-Saucedo, C., Cerna, J. F., Villegas-Sepulveda, N., Thompson, R., Velazquez, F. R., Torres, J. & Estrada-Garcia, T. (2003). Single multiplex polymerase chain reaction to detect diverse loci associated with diarrheagenic *Escherichia coli*. Emerg. Infect. Dis. 9(1), 127.

Mahmoudi-Aznaveh, A., Bakhshi, B., Najar-Peerayeh, S., Kazemnejad, A., Rafieepour, Z., Rahbar, M. & Abbaspour, S. (2013). Commensal *E. coli* as an Important Reservoir of Resistance encoding Genetic elements. Int. J. Enteric. Pathog. 1(2), 44–48.

Mao, B. H., Chang, Y. F., Scaria, J., Chang, C. C., Chou, L. W., Tien, N. et al. (2012). Identification of *Escherichia coli* genes associated with urinary tract infections. J. Clinical Microbiol. 50(2), 449–456.

Maragkoudakis, P.A., Zoumpopoulou, G., Miaris, C., Kalantzopoulos, G., Pot, B. & Tsakalidou, E. (2006). Probiotic potential of *Lactobacillus* strains isolated from dairy products. Int. Dairy J. 16:189–199.

Masomian, M., Rahman, R. N. Z. R. A., Salleh, A. B. & Basri, M. (2016). Analysis of comparative sequence and genomic data to verify phylogenetic relationship and explore a new subfamily of bacterial lipases. PLOS One 11(3), e0149851.

Matin, M. A., Islam, M. A. & Khatun, M. M. (2017). Prevalence of colibacillosis in chickens in greater Mymensingh district of Bangladesh. Vet. World 10(1), 29.

Miles, T. D., McLaughlin, W. & Brown, P. D. (2006). Antimicrobial resistance of *Escherichia coli* isolates from broiler chickens and humans. BMC Vet. Res. 2(1), 7.

Mokady, D., Gophna, U. & Ron, E. Z. (2005). Virulence factors of septicemic *Escherichia coli* strains. Int. J. Med. Microbiol. 295(6-7), 455–462.

Momtaz, S. & Hossain, M. A. (2018). Occurrence of Pathogenic and Multidrug Resistant *Salmonella* spp. Biores. Comm. 4(2), 506–515.

Mora, A., Viso, S., López, C., Alonso, M. P., García-Garrote, F., Dabhi, G. & Blanco, J. E. (2013). Poultry as reservoir for extraintestinal pathogenic *Escherichia coli* O45: K1: H7-B2-ST95 in humans. Vet. Microbiol. 167(3-4), 506–512.

Moreno, E., Prats, G., Sabaté, M., Pérez, T., Johnson, J. R. & Andreu, A. (2006). Quinolone, fluoroquinolone and trimethoprim/sulfamethoxazole resistance in relation to virulence determinants and phylogenetic background among uropathogenic *Escherichia coli*. J. Antimicrob. Chemother. 57(2), 204–211.

Moulin-Schouleur, M., Répérant, M., Laurent, S., Brée, A., Mignon-Grasteau, S., Germon, P. & Schouler, C. (2007). Extraintestinal pathogenic *Escherichia coli* strains of avian and human origin: link between phylogenetic relationships and common virulence patterns. J. Clin. Microbiol. 45(10), 3366–3376.

Murray, J., Muruko, T., Gill, C. I., Kearney, M. P., Farren, D., Scott, M. G. & Ternan, N. G. (2017). Evaluation of bactericidal and anti-biofilm properties of a novel surface-active organosilane biocide against healthcare associated pathogens and *Pseudomonas aeruginosa* biolfilm. PLoS One 12(8), 182624.

Müstak, H. K., Günaydin, E., Kaya, İ. B., Salar, M. Ö., Babacan, O. et al. (2015). Phylo-typing of clinical *Escherichia coli* isolates originating from bovine mastitis and canine pyometra and urinary tract infection by means of quadruplex PCR. Vet. Quarterly 35(4), 194–199.

Osman, K. M., Kappell, A. D., Elhadidy, M., ElMougy, F., El-Ghany, W. A. A., Orabi, A. & Hessain, A. M. (2018). Poultry hatcheries as potential reservoirs for antimicrobial-resistant *Escherichia coli*: a risk to public health and food safety. Sci. Rep. 8(1), 5859.

Pasquali, F., Lucchi, A., Braggio, S., Giovanardi, D., Franchini, A., Stonfer, M. & Manfreda, G. (2015). Genetic diversity of *Escherichia coli* isolates of animal and environmental origins from an integrated poultry production chain. Vet. Microbiol. 178(3-4), 230–237.

Prescott, J. F., Baggot, J. D. & Walker, R. D. (2000). Antimicrobial therapy in veterinary medicine. 3. USA: Iowa State Press, Ames, Iowa.

Rakhi, N. N., Alam, A. R. U., Sultana, M., Rahaman, M. M. & Hossain, M. A. (2019). Diversity of carbapenemases in clinical isolates: The emergence of blaVIM-5 in Bangladesh. J. Infect. Chem. 25(6), 444–451.

Ramadan, H., Awad, A., & Ateya, A. (2016). Detection of phenotypes, virulence genes and phylotypes of avian pathogenic and human diarrheagenic *Escherichia coli* in Egypt. J. Infect. Dev. Ctries. 10(6), 584LJ591.

Reichhardt, C., & Cegelski, L. (2018). The Congo red derivative FSB binds to curli amyloid fibers and specifically stains curliated *E. coli*. PLOS One 13(8), e0203226, https://doi.org/10.1371/journal.pone.0203226.

Reichhardt, C., Jacobson, A. N., Maher, M. C., Uang, J., McCrate, O. A., Eckart, M. et al. (2015) Congo Red Interactions with Curli-Producing *E. coli* and Native Curli Amyloid Fibers. PLOS One 10(10), e0140388, https://doi.org/10.1371/journal.pone.0140388.

Reza, S., Akond, M.A., Hassan, S.M., Alam, S. & Shirin, M. (2009). Antibiotic Resistance of *Escherichia coli* Isolated from Poultry and Poultry Environment of Bangladesh. Am. J. Environ. Sci. 5, 47–52.

Römling, U. & Balsalobre, C. (2012). Biofilm infections, their resilience to therapy and innovative treatment strategies. J. Int. Med. 272(6), 541–561.

Saitou, N. & Nei, M. (1987). The neighbor-joining method: a new method for reconstructing phylogenetic trees. Mol. Biol. Evol. 4(4), 406–425.

Salipante, S. J., Roach, D. J., Kitzman, J. O., Snyder, M. W., Stackhouse, B., Butler-Wu, S. M. et al. (2015). Large-scale genomic sequencing of extraintestinal pathogenic *Escherichia coli* strains. Genome Res. 25(1), 119–128.

Sarba, E. J., Kelbesa, K. A., Bayu, M. D., Gebremedhin, E. Z., Borena, B. M. & Teshale, A. (2019). Identification and antimicrobial susceptibility profile of *Escherichia coli* isolated from backyard chicken in and around ambo, Central Ethiopia. BMC Vet. Res. 15(1), 85.

Sarker, M. S., Mannan, M. S., Ali, M. Y., Bayzid, M., Ahad, A. & Bupasha, Z. B. (2019). Antibiotic resistance of *Escherichia coli* isolated from broilers sold at live bird markets in Chattogram, Bangladesh. J. Adv. Vet. Anim. Res. 6(3), 272.

Saud, B., Paudel, G., Khichaju, S., Bajracharya, D., Dhungana, G., Awasthi, M. S. & Shrestha, V. (2019). Multidrug-Resistant Bacteria from Raw Meat of Buffalo and Chicken, Nepal. Vet. Med. Int. 7960268.

Sekhar, M. S., Sharif, N. M., Rao, T. S. & Metta, M. (2017). Genotyping of virulent *Escherichia coli* obtained from poultry and poultry farm workers using enterobacterial repetitive intergenic consensus-polymerase chain reaction. Vet. World. 10(11), 1292–1296.

Sgariglia, E., Mandolini, N. A., Napoleoni, M., Medici, L., Fraticelli, R., Conquista, M. & Sargenti, M. (2019). Antibiotic resistance pattern and virulence genes in avian pathogenic *Escherichia coli* (APEC) from different breeding systems. Vet. Italiana 55(1), 27–33.

Smith, J. L., Fratamico, P. M. & Gunther, N. W. (2007). Extraintestinal pathogenic *Escherichia coli*. Foodborne Pathog. Dis. 4(2), 134–163.

Solà-Ginés, M., Cameron-Veas, K., Badiola, I., Dolz, R., Majó, N., Dahbi, G. et al. (2015). Diversity of Multi-Drug Resistant Avian Pathogenic Escherichia coli (APEC) Causing Outbreaks of Colibacillosis in Broilers during 2012 in Spain. PLOS One 10(11), e0143191, https://doi.org/10.1371/journal.pone.0143191.

Stebbins, M.B., Berkhoff, H.A., Corbett, W.T. (1992). Epidemiological studies of Congo Red E. coli in broiler chicken. Can. J. Vet. Res. 56, 202–205.

Subedi, M., Luitel, H., Devkota, B. et al. (2018). Antibiotic resistance pattern and virulence genes content in avian pathogenic *Escherichia coli* (APEC) from broiler chickens in Chitwan, Nepal. BMC Vet. Res. 14, 113.

Sultana, A., Saha, O., Siddiqui, A. R., Saha, A., Hussain, M. S. & Islam, T. (2018). Molecular Detection of Multidrug Resistance Pathogenic Bacteria from Protective Materials Used By Healthcare Workers (HCW); Bangladesh Scenario. J. Appl. Sci. 18(1), 48–55.

Tello, A., Austin, B. & Telfer, T. C. (2012). Selective pressure of antibiotic pollution on bacteria of importance to public health. Env. Health Pers. 120(8), 1100–1106.

Tikoo, A., Tripathi, A. K., Verma, S. C., Agrawal, N. & Nath, G. (2001). Application of PCR fingerprinting techniques for identification and discrimination of *Salmonella* isolates. Current Sci. 1049–1052.

Toma, C., Lu, Y., Higa, N., Nakasone, N., Chinen, I., Baschkier, A. & Iwanaga, M. (2003). Multiplex PCR assay for identification of human diarrheagenic *Escherichia coli*. J. Clinical Microbiol. 41(6), 2669–2671.

Tsai, Y., Palmer, C. J. & Sangermano, L. R. (1993). Detection of *Escherichia coli* in sewage sludge by polymerase chain reaction. Appl. Environ. Microbiol. 59(2): 353–357.

Wang, S., Liu, X., Xu, X., Yang, D., Wang, D., Han, X. & Yu, S. (2016). *Escherichia coli* Type III Secretion System 2 ATPase EivC Is Involved in the Motility and Virulence of Avian Pathogenic *Escherichia coli*. Front. Microbiol. 7(1387).

Wang, X. M., Liao, X. P., Zhang, W. J., Jiang, H. X., Sun, J., Zhang, M. J. et al. (2010). Prevalence of serogroups, virulence genotypes, antimicrobial resistance, and phylogenetic background of avian pathogenic *Escherichia coli* in south of China. Foodborne Pathog. Dis. 7(9), 1099–1106.

Xu, Y., Bai, X., Jin, Y., Hu, B., Wang, H., Sun, H. et al. (2017). High prevalence of virulence genes in specific genotypes of atypical enteropathogenic *Escherichia coli*. Front. Cell. Infect. Microbiol. 7, 109, https://doi.org/10.3389/fcimb.2017.00109.

Yamamoto, S., Terai, A., Yuri, K., Kurazono, H., Takeda, Y. & Yoshida, O. (1995). Detection of urovirulence factors in *Escherichia coli* by multiplex polymerase chain reaction. FEMS Immun. Med. Microbiol. 12(2), 85–90.

Yoon, M. Y., Min, K. B., Lee, K.-M., Yoon, Y., Kim, Y., Oh, Y. T. et al. (2016). A single gene of a commensal microbe affects host susceptibility to enteric infection. Nat. Commun. 7, 11606.

Younis, G., Awad, A. & Mohamed, N. (2017). Phenotypic and genotypic characterization of antimicrobial susceptibility of avian pathogenic *Escherichia coli* isolated from broiler chickens. Vet. World 10(10), 1167–1172.

Zahid, A. A., AL-Mossawei, M. T.,Mahmood, A. B. (2016). In vitro and In vivo Pathogenicity tests of Local Isolates APEC from Naturally Infected Broiler in Baghdad. Int. J. Adv. Res. Biol. Sci. 3(3), 89–100.

Zhao, L., Chen, X., Zhu, X., et al. (2009). Prevalence of virulence factors and antimicrobial resistance of uropathogenic *Escherichia coli* in Jiangsu province (China). Urology 74(3), 702□707, doi: 10.1016/j.urology.2009.01.042.

