## Supplementary Table 1 for "Multidrug-Resistant Avian Pathogenic *Escherichia coli* Strains and Association of Their Virulence Genes in Bangladesh"

**Supplementary Table 1:** Sequence of oligonucleotide primers of different target genes used in this study to detect pathogenic *Escherichia coli* strains.

| **Target gene** | **DNA sequence (5′→3′)** | **Amplified product (bp)** | **References** |
| --- | --- | --- | --- |
| *ial* | GGTATGATGATGATGAGTCCA  GGAGGCCAACAATTATTTCC | 650 | Lopez-Saucedo *et al.* (2003) |
| *bfpA* | AATGGTGCTTGCGCTTGCTGC  GCCGCTTTATCCAACCTGGTA | 334 |  |
| *Stx1* | CTGGATTTAATGTCGCATAGTG  AGAACGCCCACTGAGATCATC | 150 |  |
| *Stx2* | GGCACTGTCTGAAACTGCTCC  TCGCCAGTTATCTGACATTCTG | 255 |  |
| *eaeA* | GACCCGGCACAAGCATAAGC  CCACCTGCAGCAACAAGAGG | 384 |  |
| *lt* | GGC GAC AGA TTA TAC CGT GC  CGG TCT CTA TAT TCC CTG TT | 450 | Houser *et al.*(2008) |
| *aggR* | GTATACACAAAAGAAGGAAGC  ACAGAATCGTCAGCATCAGC | 254 | Toma *et al*. (2003) |
| *uidA* | AAAACGGCAAGAAAAAG CAG  ACGCGTGGTTACAGTCTT GCG | 147 | Bej *et al*.(1991) |
| *cjrC* | AAACCTCAGCGCAAAATCGT  AGGCTTCAGGAATGGGTTCA | 518 | Mao *et al*.(2012) |
| *fimH* | GTGCCAATTCCTCTTACCGTT  TGGAATAATCGTACCGTTGCG | 164 | Hojati *et al.*(2015) |
| *crl* | TTTCGATTGTCTGGCTGTAT  CTTCAGATTCAGCGTCGTC | 250 | Maurer *et al*.(1998) |
| *papC* | GACGGCTGTACTGCAGGGTGTGGCG  ATATCCTTTCTGCAGGGATGCAATA | 328 | Yamamoto *et al*.(1995) |
| *hly*A | GCATCATCAAGCGTACGTTCC  AATGAGCCAAGCTGGTTAAGCT | 534 | Srivani et al. (2017) |
| *ChuA* | GACGAACCAACGGTCAGGAT  TGCCGCCAGTACCAAAGACA | 279 | Clermont *et al*.(2013) |
| *Yja A* | TGAAGTGTCAGGAGACGCTG  ATGGAGAATGCGTTCCTCAAC | 211 |  |
| TspE4C2 | GAGTAATGTCGGGGCATTCA  CGCGCCAACAAAGTATTACG | 152 |  |
| *arp*A | AACGCTATTCGCCAGCTTGC  TCTCCCCATACCGTACGCTA | 400 |  |
| *arpA* | GATTCCATCTTGTCAAAATATGCC  GAAAAGAAAAAGAATTCCCAAGAG | 301 |  |
| *trpA* | AGTTTTATGCCCAGTGCGAG  TCTGCGCCGGTCACGCCC | 219 |  |

**arp*A for phylotype E and *trp*A for phylotype C
