## Supplementary Table 2 for "Multidrug-Resistant Avian Pathogenic *Escherichia coli* Strains and Association of Their Virulence Genes in Bangladesh"

**Supplementary Table 2:** Relative comparison among isolated Pathogenic *E. coli* from different poultry farms of three sampling locations.

| Farm | Sources | Isolates ID | Phylotype | Congo Red Assay | Pathogenic genes Number | Biofilm Formation | Antibiotics pattern |
| --- | --- | --- | --- | --- | --- | --- | --- |
| Farm 1 | DR | DH21 | A1 | ++ | ND | 2 | ND |
|  |  | DH22 | A1 | ++ | ND | 3 | ND |
|  |  | DH25 | B2(B2_3_) | +++ | *uid*A*crlfim*H*ialpap*C*cjr*C | 1 | DoTeFCIPNAGNCS |
|  |  | DH26 | D2 | + | *uid*A*crlfim*H*ialpap*C | 0 | ND |
|  |  | DH27 | A1 | +++ | ND | 0 | ND |
|  |  | DH28 | D2 | ++ | *uidAcrlfim*H*ialpap*C*cjr*C | 0 | ND |
|  |  | DH52 | B2(B2_3_) | ++ | ND | 0 | AMPDoTeFCIPGNCS |
|  |  | DH53* | B1 | +++ | *uid*A*crlfim*H*ial* | 4 | AMPDoTeCIPNACSATM |
|  |  | DH55 | B2(B2_3_) | ++ | ND | 2 | AMPDoFCIPNAIMPC |
|  |  | DH57 | D2 | ++ | *uid*A*crlfim*H*ialpapCcjrC* | 0 | DoTeFNAFoxGNCS |
|  |  | DH59* | D2 | +++ | *uid*A*crlfim*H*ialpap*C*cjr*C | 2 | DoTeFCIPNAFoxIMPS |
|  |  | DH60 | B1 | +++ | *uid*A*crlfim*H | 0 | ND |
|  | CS | DH10 | B2(B2_3_) | ++ | *uid*A*crlfim*H*ial* | 3 | AMPTeFNAGNNA |
|  |  | DH23 | A1 | +++ | *uid*A*crlial* | 4 | ND |
|  |  | DH24 | A1 | ++ | *uid*A*crlial* | 0 | DoTeFNACS |
|  |  | DH31 | B2(B2_3_) | ++ | ND | 0 | DoTeFNASATM |
|  |  | DH32 | A1 | + | ND | 0 | AMPTeFNAFoxS |
|  |  | DH33 | A1 | ++ | *uid*A*crlial* | 0 | AMPDoTeFoxSATM |
|  |  | DH34 | A1 | +++ | *uid*A*crlfim*H*ial* | 0 | ND |
|  |  | DH35 | A1 | ++ | ND | 0 | AMPDoFCIPFoxC |
|  |  | DH42 | B1 | ++ | ND | 2 | AMPDoTeCIPFoxCATM |
|  |  | DH49 | B2(B2_3_) | +++ | *uid*A*crlfim*H*ial* | 3 | AMPTeFCIPNAFoxIMPC |
|  |  | DH67 | B2(B2_3_) | + | ND | 0 | DoTeFNAS |
|  |  | DH69* | A1 | +++ | *uid*A*crlfim*H*ialpap*C | 3 | AMPDoPBCIPNAFoxIMP |
|  | F | DH1 | A1 | ++ | *uid*A*fim*H*ialpap*C | 4 | DoTeFC |
|  |  | DH8* | B2(B2_3_) | ++ | *uid*A*crlfim*H*ial* | 3 | AMPDoTeFoxIMPCATM |
|  |  | DH48 | B2(B2_3_) | ++ | *uid*A*crlfim*H*ial* | 1 | ND |
|  |  | DH66* | B2(B2_2_) | +++ | *uid*A*crlfim*H*ial* | 2 | AMPDoTeFCATM |
|  | W | DH29 | A1 | ++ | *uid*A*crlfim*H*ial* | 2 | FCIPNAFoxGNATM |
|  |  | DH43 | B2(B2_3_) | +++ | *uid*A*crlfim*H*ial* | 2 | ND |
|  |  | DH44 | A1 | ++ | *uid*A*crlfim*H*ial* | 0 | AMPDoNAFoxIMPCATM |
|  |  | DH50 | B2(B2_3_) | ++ | ND | 2 | TeFNACSATM |
|  |  | DH51 | D2 | + | *uid*A*crlfim*H*ial* | 3 | AMPDoTeFNAIMPC |
|  | H | DH20# | A1 | ++ | ND | 1 | DoTeFNAS |
| Farm 2 | DR | DH86* | B2(B2_3_) | +++ | *uid*A*crlfim*H*ial* | 3 | AMPDoTeFCIPNAFoxS |
|  |  | DH87 | B2(B2_3_) | +++ | *uid*A*crlfim*H*ial* | 0 | TeFFoxCSATM |
|  |  | DH89 | D2 | +++ | *uid*A*crlfim*H*ial* | 1 | AMPDoTeFPBNAFoxIMP |
|  |  | DH90 | B1 | +++ | *uid*A*crlial* | 0 | ND |
|  |  | DH93# | A1 | +++ | *uid*A*crlfim*H*ialpap*C | 0 | AMPTePBNAFoxGNCS |
|  |  | DH95 | A1 | ++ | ND | 0 | ND |
|  |  | DH96 | B2(B2_3_) | +++ | *uid*A*crlfim*H*ialcjr*C | 1 | AMPDoFFoxCATM |
|  |  | DH97 | B2(B2_3_) | +++ | *uid*A*crlfim*H*ial* | 3 | DoTeFoxSATM |
|  | CS | DH73# | B1 | +++ | *uid*A*crlfim*H | 0 | AMPTeFCIPNAGNCATM |
|  |  | DH74 | B1 | +++ | *uid*A*crlfim*H | 0 | ND |
|  |  | DH75# | B1 | +++ | *uid*A*crlfim*H*ial* | 0 | AMPDoTeFCIPNAGNS |
|  | F | DH72 | B2(B2_3_) | ++ | ND | 0 | DoTeFFoxCSATM |
|  | W | DH71 | D2 | + | ND | 0 | AMPDoTeCIPNAFoxS |
|  |  | DH91 | B2(B2_3_) | + | *uid*A*crlfim*H*ial* | 0 | ND |
|  |  | DH94 | A1 | + | *ND* | 4 | ND |
|  |  | DH100* | B2(B2_3_) | +++ | *uid*A*crlfim*H*ialcjr*C | 1 | AMPDoTeFGNS |
|  | H | DH82 | D2 | ++ | *uid*A*crlfim*H*ial cjr*C | 3 | ND |
|  |  | DH83* | A1 | +++ | *uid*A*crlfim*H | 0 | AMPDoTeFCIPNAFoxIMPGNS |
|  |  | DH85 | B2(B2_3_) | +++ | *uid*A*crlfimHialpap*C | 0 | ND |
|  | IO(L) | DH76 | B1 | ++ | ND | 0 | AMPDoTeFCIPNAFox |
|  |  | DH80* | B2(B2_2_) | +++ | *uid*A*crlfim*H*ialpapC* | 0 | AMPDoPBNAFoxCATM |
|  |  | DH103 | B2(B2_3_) | +++ | *uid*A*crlfim*H*ial* | 0 | ND |
|  |  | DH106* | D2 | +++ | *uid*A*crlfim*H*ialpap*C*cjr*C | 1 | AMPDoTeCIPNAGNCS |
|  |  | DH108 | B1 | ++ | *uid*A*crlfim*H*ial* | 4 | AMPDoTeNAIMPGN |
|  |  | DH109 | B2(B2_3_) | +++ | *uid*A*crlfim*H*ialpap*C | 0 | ND |
| Farm 3 | DR | DH152 | B2(B2_3_) | ++ | *ND* | 0 | AMPDoTeNAGNATM |
|  |  | DH162 | A1 | +++ | *uid*A*pap*C*cjr*C | 0 | ND |
|  |  | DH185 | B1 | + | ND | 0 | ND |
|  |  | DH170 | A1 | + | ND | 2 | AMPDoFNAGNS |
|  |  | DH116* | B2(B2_2_) | +++ | *uid*A*crlfim*H*ial* | 3 | AMPDoTeFCIPNAIMPFoxGNCSATM |
|  | W | DH141* | B2(B2_2_) | +++ | *uid*A*crlfim*H*ialpap*C | 0 | AMPDoTeCIPNAIMPC |
|  | H | DH168 | B2(B2_3_) | ++ | *uid*A*crlfim*H*ial* | 2 | AMPDoTeFNAFoxATM |
|  |  | DH179 | A1 | ++ | *uid*A*crl pap*C | 4 | ND |
|  | ES | DH166 | B1 | +++ | *uid*A*crlfim*H*ial* | 3 | ND |
|  |  | DH187 | A1 | + | ND | 0 | AMPDoFCIPNAFoxC |
|  | IO(L) | DH144 | B2(B2_3_) | + | ND | 0 | NS |
|  |  | DH145 | B2(B2_3_) | +++ | *uid*A*crlfim*H*ialcjr*C | 3 | AMPDoTeFCIPNAIMPC |
|  |  | DH149 | B2(B2_3_) | +++ | *uid*A*crlfim*H*ialpap*C | 4 | ND |
|  |  | DH173 | B2(B2_3_) | +++ | *uid*A*crlfim*H*ial* | 1 | ND |
|  |  | DH175 | B2(B2_3_) | +++ | *uid*A*crlfim*H*ial* | 4 | AMPDoTeFCIPNAGNSATM |
|  |  | DH191 | B2(B2_2_) | +++ | ND | 0 | AMPTeFNAGNIMPCATM |
|  |  | DH196* | A1 | ++ | ND | 0 | AMPTeFNAS |
| Farm 4 | DR | RN3(2)* | B1 | +++ | *uid*A*crlfim*H*ial* | 1 | AMPDoTeCIPGNCSATM |
|  |  | RN07# | A1 | +++ | *uid*A*crlfim*H*ial* | 2 | AMPDoTeFNA |
|  |  | RN08 | A1 | + | ND | 0 | ND |
|  |  | RN09 | A1 | ++ | *uid*A*fim*H*ial* | 0 | ND |
|  |  | RN10* | A1 | +++ | *uid*A*crlfim*H*ialpap*C | 1 | AMPTeNAGNS |
|  |  | RN14* | B2(B2_2_) | ++ | *uid*A*crlfim*H*ial* | 1 | AMPDoTeFCIPIMPC |
|  |  | RN22 | B2(B2_2_) | +++ | ND | 0 | AMPDoTeCIPIMPC |
|  |  | RN33 | B1 | +++ | ND | 2 | ND |
|  |  | RN34 | A1 | +++ | *uid*A*fim*H*ialcjr*C | 4 | AMPFNAGNCS |
|  |  | RN35# | B2(B2_2_) | ++ | ND | 3 | AMPDoTeFNASCATM |
|  |  | RN62 | D2 | + | *uid*A*crlfim*H*ial* | 0 | DoFIMP |
|  |  | RN63 | B2(B2_3_) | +++ | *uid*A*crlfim*H*ialpap*C | 1 | ND |
|  |  | RN64 | A1 | +++ | *uid*A*crlfim*H*ial* | 0 | AMPDoTeF |
|  |  | RN65 | A1 | +++ | *uid*A*crlfim*H*ial* | 0 | ND |
|  | CS | RN36 | B2(B2_3_) | ++ | *uid*A*crlfim*H*ialcjr*C | 4 | ND |
|  |  | RN38 | B2(B2_3_) | + | ND | 1 | AMPDoTeCIPNAC |
|  |  | RN39 | B2(B2_3_) | ++ | ND | 0 | AMPDoTeFFoxGN |
|  |  | RN41# | B2(B2_3_) | +++ | *uid*A*crlfim*H*ial* | 3 | DoTeCS |
|  |  | RN66# | A1 | +++ | ND | 0 | AMPDoTeCIPNAC |
|  |  | RN68 | A1 | ++ | ND | 4 | ND |
|  | F | RN42* | B2(B2_2_) | +++ | *uid*A*crlfim*H*ial cjr*C | 0 | AMPDoTeNA |
|  |  | RN43 | A1 | ++ | *uid*A*crlfim*H*ial* | 0 | ND |
|  | W | RN32 | D2 | + | *uid*A*crlfim*H*ialpap*C*cjr*C | 1 | AMPDoTeCIPNAFoxS |
|  | H | RN16 | A1 | +++ | *uid*A*crlfim*H | 0 | AMPDoFCIPFoxATM |
|  |  | RN18 | B2(B2_3_) | +++ | *uid*A*crlfim*H*ialpap*C | 4 | AMPDoTeFFoxCS |
|  |  | RN19 | B2(B2_2_) | ++ | *uid*A*crlfim*H*ial* | 4 | TeFCIPIMPCATM |
|  |  | RN29 | D2 | +++ | *uid*A*crlfim*H*ialpapCcjrC* | 0 | AMPDoTeNAFoxIMP |
|  |  | RN30 | B2(B2_3_) | ++ | *uid*A*crlfim*H*ial* | 2 | TeFCIPFoxS |
|  |  | RN31 | B2(B2_3_) | ++ | *uid*A*crlfim*H*ial* | 0 | DoTeNAFoxGN |
|  | IO(L)) | RN20 | B2(B2_3_) | + | ND | 0 | ND |
|  |  | RN46 | B2(B2_3_) | + | ND | 4 | AMPDoFCATM |
|  |  | RN47 | A1 | ++ | *uid*A*crlfim*H*ial* | 0 | ND |
|  |  | RN48 | B2(B2_3_) | +++ | *uidAcrlfimHialcjrC* | 0 | AMPTeFPBNAGNATM |
|  |  | RN50* | B2(B2_2_) | +++ | *uid*A*crlfim*H*ial cjr*C | 0 | AMPDoTeFCIP |
|  |  | RN51* | B2(B2_3_) | +++ | *uid*A*crlfim*H*ial* | 0 | DoTeCIPNAFoxCATM |
|  |  | RN52 | B2(B2_3_) | ++ | ND | 2 | ND |
|  |  | RN53 | A1 | + | ND | 0 | ND |
|  |  | RN54 | A1 | +++ | *uid*A*ialpap*C | 2 | AMPFPB |
|  |  | RN55 | A1 | +++ | *uid*A*ialpap*C | 0 | AMPTeCIPFoxIMPS |
|  |  | RN58 | B2(B2_3_) | ++ | ND | 0 | AMPDoTeFCIP |
|  |  | RN59* | D2 | +++ | *uid*A*crlfim*H*ialpap*C*cjr*C | 2 | DoTeFCIPGNCS |
|  |  | RN60 | A1 | ++ | ND | 0 | ND |
| Farm 5 | DR | RN93* | B1 | +++ | *uid*A*crlfim*H*ial* | 4 | AMPDoTeFNAFoxS |
|  |  | RN105 | B2(B2_2_) | ++ | *uid*A*crlfim*Hi*alpap*C | 0 | ND |
|  |  | RN106 | B2(B2_3_) | ++ | ND | 3 | DoTeCIPNAC |
|  |  | RN110 | A1 | +++ | *uid*A*crlfim*H*ial* | 0 | DoTeCIPNAS |
|  |  | RN111 | A1 | ++ | *uid*A*crlfim*H*ial* | 0 | AMPDoTeNAFoxGNC |
|  |  | RN112 | A1 | ++ | *uid*A*crlfim*H*ial* | 0 | AMPTeFNAFoxGN |
|  |  | RN113 | B2(B2_2_) | +++ | *uid*A*crlfim*H*ialcjr*C | 0 | ND |
|  |  | RN114 | B2(B2_2_) | + | ND | 1 | ND |
|  |  | RN117* | B2(B2_3_) | ++ | *uid*A*crlfim*H*ialcjr*C | 2 | AMPDoTeFNASATM |
|  |  | RN118# | A1 | +++ | *uid*A*crlfim*H*ialcjr*C | 0 | AMPDoTeCIPNAIMPC |
|  |  | RN122* | A1 | +++ | *uid*A*crlfim*H | 1 | DoTeFC |
|  |  | RN133 | B2(B2_3_) | +++ | *uid*A*crlfim*H*ialpap*C | 0 | AMPDoTePBCIPGNC |
|  |  | RN135 | B2(B2_3_) | ++ | ND | 0 | ND |
|  |  | RN138 | B2(B2_3_) | +++ | *uid*A*crlfim*H*ial cjr*C | 0 | AMPDoTeCIPFoxCATM |
|  |  | RN139 | B2(B2_3_) | ++ | *uid*A*crlfim*H*ial cjr*C | 0 | DoTeFGN |
|  | CS | RN99 | B2(B2_3_) | + | ND | 3 | ND |
|  |  | RN100 | B2(B2_2_) | ++ | *uid*A*crlfim*H*ialpap*C | 0 | ND |
|  |  | RN101* | D2 | +++ | *uid*A*crlfim*H*ialpap*C*cjr*C | 1 | ND |
|  |  | RN102 | D2 | ++ | *uid*A*crlfim*H*ial* | 0 | ND |
|  |  | RN119 | A1 | ++ | ND | 0 | AMPTeFS |
|  | W | RN123 | A1 | ++ | *uid*A*crlfim*H*ial* | 0 | ND |
|  |  | RN128 | A1 | +++ | *uid*A*crlfim*H*ial* | 0 | ND |
|  |  | RN131 | A1 | +++ | *uid*A*crlfim*H*ial* | 0 | ND |
|  |  | RN132 | A1 | ++ | ND | 0 | AMPDoFIMPS |
|  | ES | RN83* | B2(B2_3_) | ++ | *uid*A*crlfim*H*ial* | 0 | AMPDoTeFCIPFox |
|  |  | RN84 | A1 | +++ | *uid*A*crlfim*H*ial* | 0 | AMPDoFCSATM |
|  |  | RN85 | A1 | +++ | *uid*A*ialpap*C | 0 | AMPDoTeFNAATM |
|  |  | RN86 | A1 | ++ | ND | 0 | AMPDoTePBCIPFoxGN |
|  |  | RN87 | B2(B2_3_)* | +++ | *uid*A*crlfim*H*ial* | 0 | AMPTeC |
|  |  | RN88 | B2(B2_3_) | +++ | *uid*A*crlfim*H*ial* | 1 | ND |
|  | H | RN70 | D2 | ++ | *uid*A*crlfim*H*ialpap*C*cjr*C | 1 | ND |
|  |  | RN71* | D2 | +++ | *uid*A*crlfim*H*ialpap*C*cjr*C | 0 | AMPDoTeFCIPGNS |
|  |  | RN89# | A1 | +++ | *uid*A*crlfim*H*ial* | 0 | DoTeFCIP |
|  |  | RN90 | B2(B2_3_) | ++ | *uid*A*crlfim*H*ialpap*C*cjr*C | 0 | AMPFCIPFoxIMPCATM |
|  |  | RN95 | B2(B2_3_) | ++ | ND | 3 | ND |
|  |  | RN96* | B2(B2_3_) | +++ | *uid*A*crlfim*Hi*alpap*C*cj*rC | 1 | AMPTeCIPCS |
|  | IO(L) | RN97* | B2(B2_3_) | +++ | *uid*A*crlfim*H*ialpapC* | 0 | AMPDoTeCIPFox |
|  |  | RN126 | D2 | ++ | *uidAcrlfimHialpap*C | 0 | ND |
|  |  | RN98 | B2(B2_3_) | ++ | *uid*A*crlfim*H*ialpap*C | 2 | AMPDoTeFC |
| Farm 6 | DR | NR1* | B2(B2_3_) | +++ | *uid*A*crlfim*H*ialcjr*C | 1 | AMPDoTeFNAC |
|  |  | NR3 | B2(B2_3_) | +++ | ND | 0 | ND |
|  |  | NR4 | B2(B2_3_) | ++ | ND | 0 | ND |
|  |  | NR6 | D2 | +++ | *uid*A*crlfim*H*ialpap*C*cjr*C | 0 | ND |
|  |  | NR9 | B2(B2_3_) | ++ | ND | 1 | AMPDoTeNAFoxS |
|  |  | NR10# | B2(B2_3_) | +++ | *uid*A*crlfim*H*ialpap*C | 2 | AMPDoTeFPBCIPNAIMPCATM |
|  | CS | NR13 | B2(B2_3_) | + | ND | 0 | AMPDoTeNACS |
|  |  | NR14 | A1 | +++ | *uid*A*crlfim*H*ial* | 2 | AMPDoTeFCIPNACS |
|  |  | NR15 | A1 | + | ND | 3 | ND |
|  |  | NR20 | A1 | ++ | *uid*A*crlfim*H*ial* | 3 | ND |
|  | F | NR28 | B2(B2_3_) | +++ | *uid*A*crlfim*H*ial* | 0 | ND |
|  |  | NR34# | A1 | +++ | *uid*A*crlfim*H*ialcjr*C | 0 | TeFNAFoxCS |
|  | IO(L) | NR35 | B2(B2_3_) | +++ | *uid*A*crlfim*H*ial* | 2 | AMPDoTeFNAFoxIMPCS |
|  |  | NR45* | A1 | +++ | *uid*A*crlfim*H*ialcjr*C | 1 | AMPTeNAFoxSATM |
|  |  | NR46* | B2(B2_2_) | ++ | *uid*A*fim*H*pap*C*cjr*C | 2 | AMPDoTeNAFoxIMPGNATM |
|  |  | NR47 | A1 | ++ | *uid*A*crlfim*H*ial* | 2 | FCIPNACSATM |
|  |  | NR48* | D2 | +++ | *uid*A*crlfim*H*ialpap*C*cjr*C | 1 | DoTeFNACS |

DH- Dhamrai(Manikgonj), RN- Rupganj(Narayangonj), NR- Norshingdi, DR-Dropping, CS-Cloacal Samples, F- Feed, W-Feeding Water; H=Handler Swab; ES= Egg Surface Swab; IO(L)=Internal Organ(Liver); ND- Not Done; AMP=ampicillin; Te=tetracycline; Do=doxycycline; F=nitrofurantoin; Pb=polymexin CIP=ciprofloxacin; NA=nalidixic acid; Fox=cefoxitin; IMP= imipenem; GN=gentamycin; C=chloramphenicol S3=sulfonamide; AZM=azithromycin; +=Positive; 0=Not Done; 1=Not Biofilm Producer; 2=Weak biofilm producer ; 3= Moderate biofilm producer; 4=Strong biofilm producer.*-Plasmid Positive;# Plasmid Negative.
