## Supplementary material for "Multidrug-Resistant Avian Pathogenic *Escherichia coli* Strains and Association of Their Virulence Genes in Bangladesh": Main Figure Legends

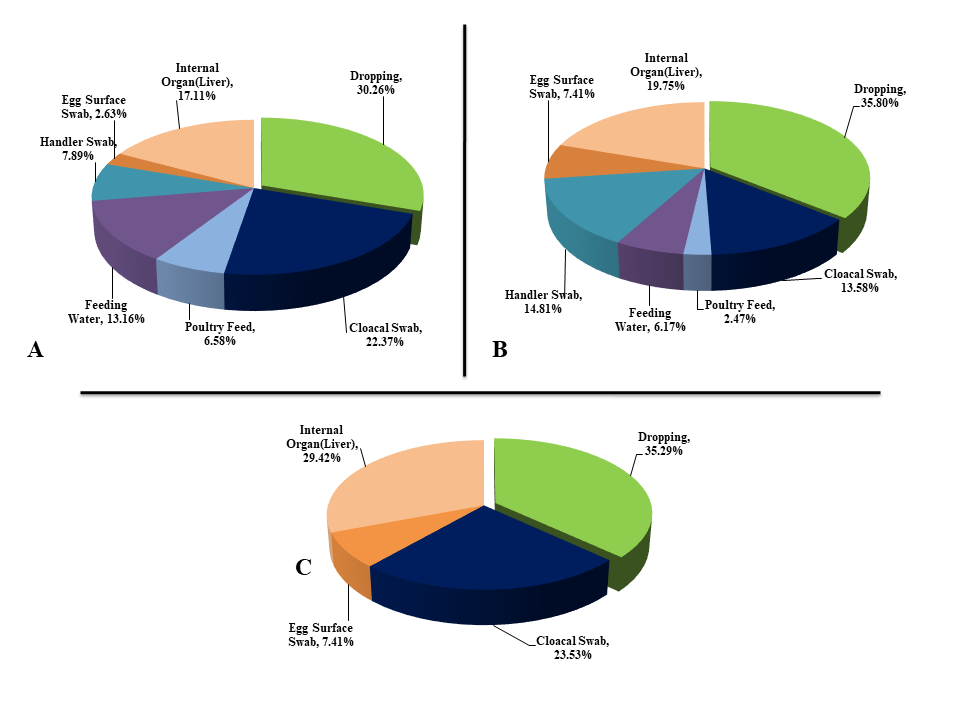

**FIGURE 1|** Occurrence of *Escherichia coli* in different types of poultry samples. Microorganisms identified in A (Dhamrai, Manikgonj); B (Rupganj, Narayangonj); and C (Monohardi, Narshingdi)) poultry samples. The Dhamrai, Rupganj and Monohardi region includes 76, 81 and 17 isolates, respectively.

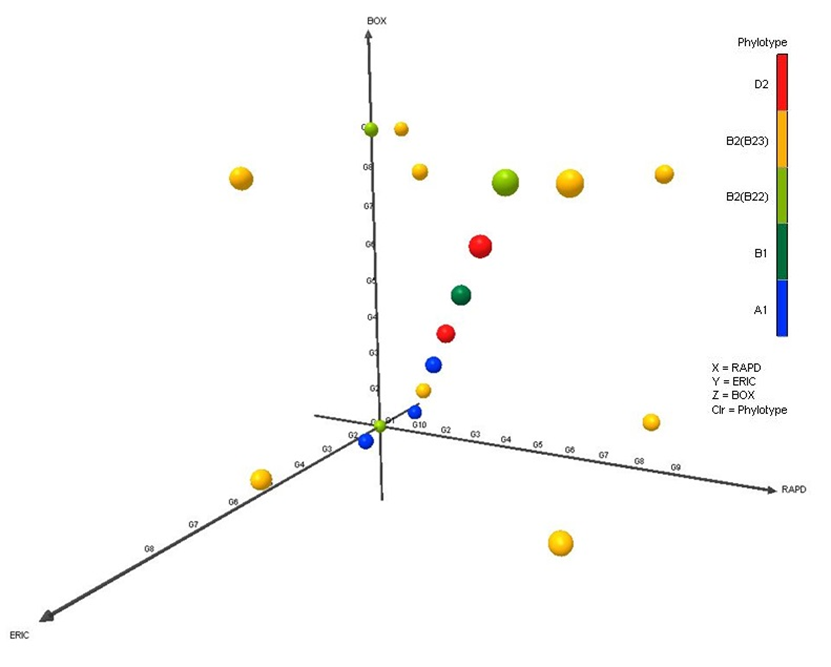

**FIGURE 2|** The diversity of APEC isolates according to various typing systems. The Principle Components Analysis (PCA) plots represent the distribution of different phylogrpoups. Orange is for phylogroup D2, blue is for A1, Yellow is for B2_3_, Green is for B2_2_ and dark red green is for B1.

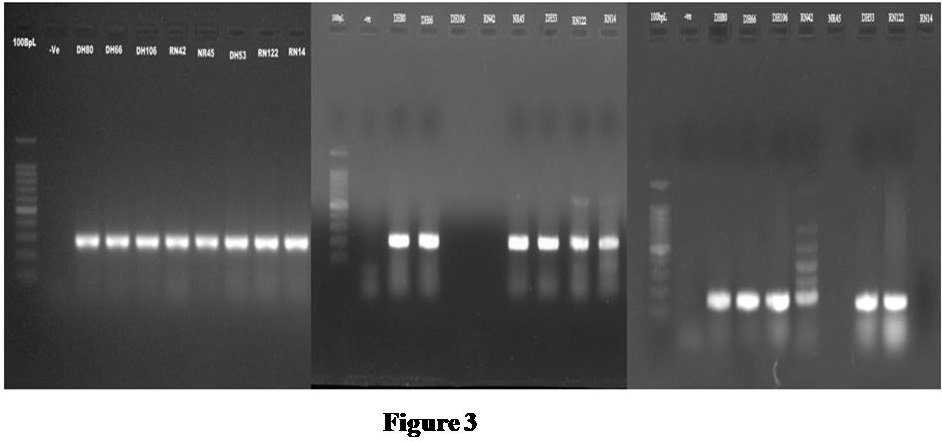

**FIGURE3 |** Representative PCR results for the detection of *Escherichia coli* phylogenetic groups among the colibacillosis cases of Bangladeshi poultry samples. A) (*chu*A: 279bp); B) (*yja*A: 211bp); C) (tspE4.C2:152bp); Here, Lane 1 is molecular ladders (100bp), Lane 2 is negative blank control and Lanes 3–10 are the strains DH80, DH66, DH106, RN42, NR45, DH53, RN122, RN14 respectively. DNA bands at the appropriate position was observed in *E.coli* strains DH66,DH80,DH53, RN122 denoted as phylogroup B2(B2_3_)(*chu*A+, *yja*A+, tspE4.C2+); *E.coli* strain DH106, RN42 denoted as phylogroup D2 (*chu*A+, *yja*A-, tspE4.C2+); *E.coli* strain NR45, RN14 denoted as phylogroup B2(B2_2_)(*chu*A+, *yja*A+, tspE4.C2-).

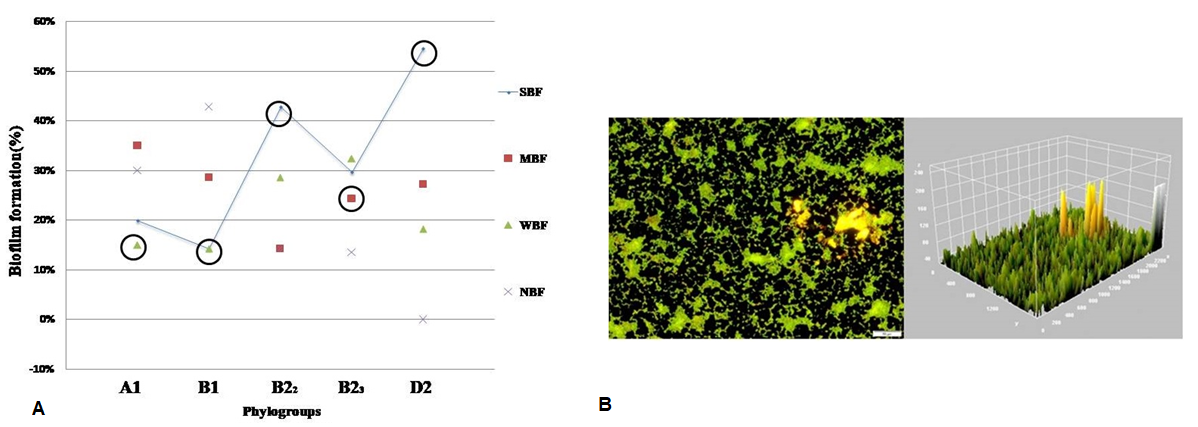

**FIGURE 4|** *Escherichia coli* biofilm development; A) Diagrammatic representation of Biofilm formation of various phylotypes. Here SBF, Strong biofilm formers; MBF, Moderate biofilm formers; WBF, Weak biofilm formers; NBF, Non biofilm formers. Solid line with circle represented the SBF ability fluctuation between the phylotypes. X axis represent the phylotypes. Y axis represented the zone of percentages of biofilm forming isolates. B) Fluoroscence microscopy images of isolate (RN3 (2)) under 20x magnification. Biofilm stained with Film tracer LIVE/DEAD biofilm viability kit. Live or active cells are fluorescent green and dead or inactive cells are fluorescent red. Surface plot of 3D volume image (center image) and cross section of 3D volume image (right side image) show the distribution of live and dead cells throughout biofilm layers.

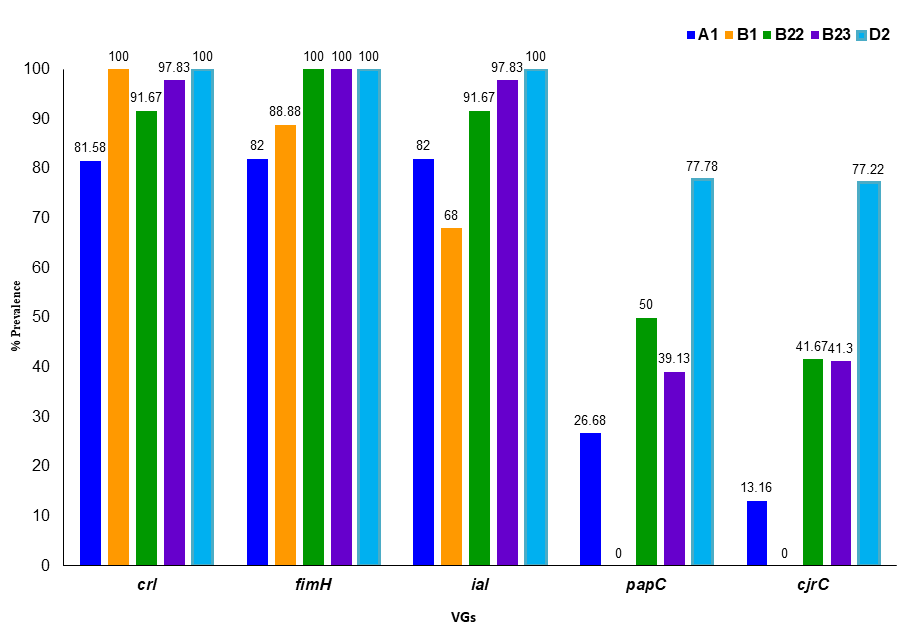

**FIGURE 5|** Prevalence of the three pathogenic genes among *E. coli* phylotypes. Here X axis represents the phylotypes. Y axis represents the prevalence (%) of the genes among the *E.coli* isolates. For phylotype A1: 1st column represents the *crl* genes; 2nd column for *fim*H; 3^rd^ Coolum *ial*; 4^th^ for *pap*C and 5^th^ for *cjr*C. This serial is true for all the others phylotypes.

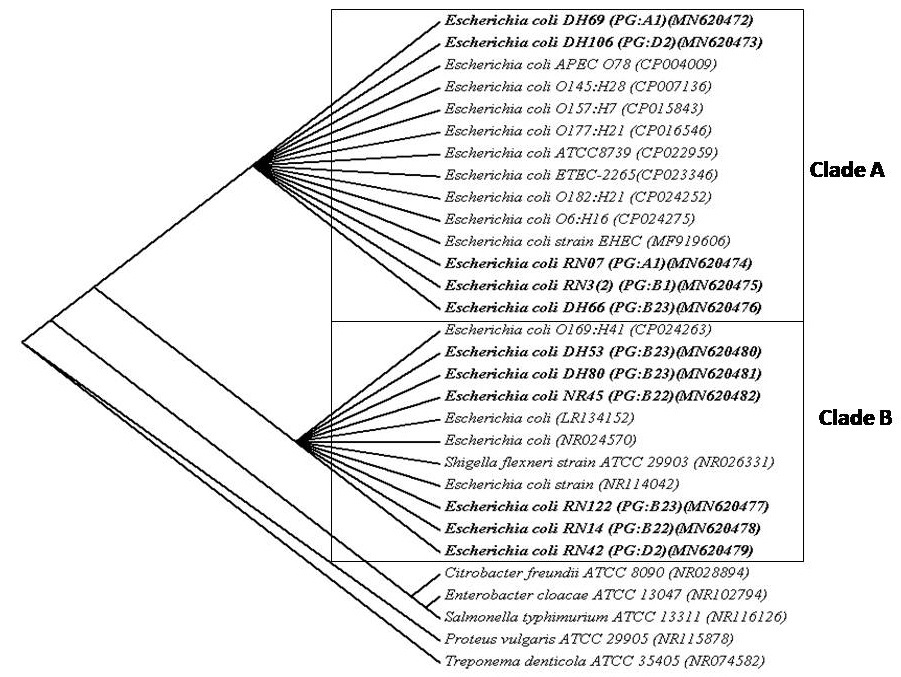

**FIGURE 6|** Phylogenetic tree predicted by the neighbor-joining method using 16S rRNA gene sequences. The evolutionary distances were computed using the Kimura 2-parameter model method and are in the units of the number of base substitutions per site. The bootstrap considered 1000 replicates. The scale bar represents the expected number of substitutions averaged over all the analyzed sites. The optimal tree with the sum of branch length = 0.35475560is shown here. *Treponema denticola* was used as out group. The length of the scale bar represents 1 nucleotide substitution per 100 positions.

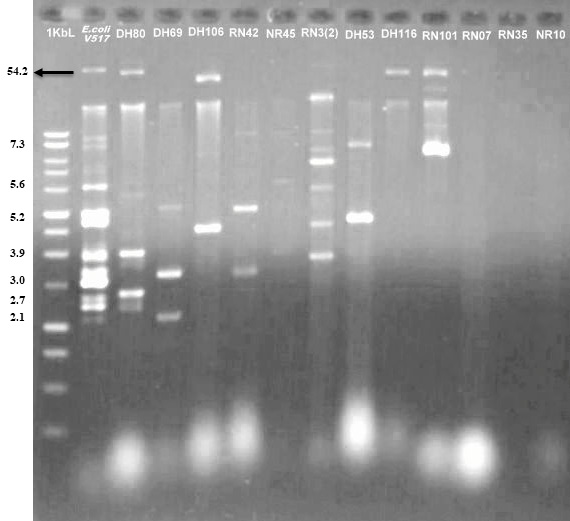

**FIGURE 7|** Phylogenetic tree predicted by the neighbor-joining method using 16S rRNA gene sequences. The evolutionary distances were computed using the Kimura 2-parameter model method and are in the units of the number of base substitutions per site. The bootstrap considered 1000 replicates. The scale bar represents the expected number of substitutions averaged over all the analyzed sites. The optimal tree with the sum of branch length = 0.35475560is shown here. *Treponema denticola* was used as out group. The length of the scale bar represents 1 nucleotide substitution per 100 positions.

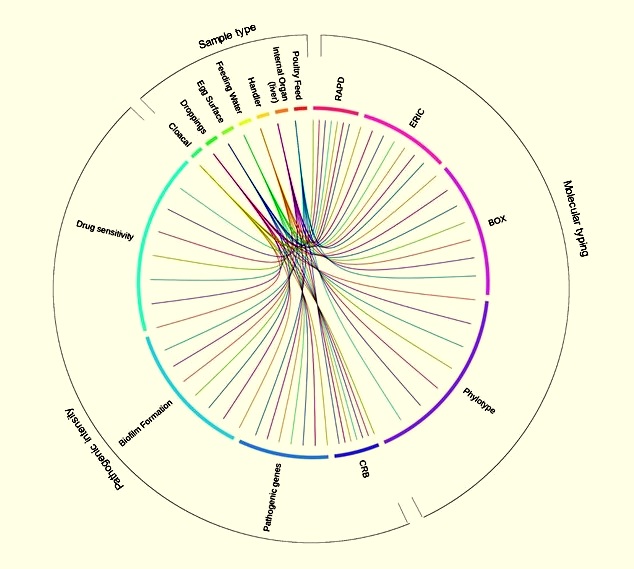

**FIGURE 8 |** Circosplot representation of association of different sample types with molecular typing and pathogenic intensity characterization of *E. coli*. The association showed significant (*p*=0.032) correlation. The frequency of occurrence of different features such as sample categories, typing and pathogenicity patterns is depicted in the outer ring. The inner ring of Circos plot depicts the correlationbetween the sample categories (cloacal, droppings, egg surface, feeding water, handler, internal organ, feed), and molecular typing (RAPD, ERIC, BOX, phylotype) and pathogenic intensities (CRB, pathogenic genes, biofilm formation, drug sensitivity). Each factor has been assigned to a color. The arc originates from sample types and terminates at typing and pathogenic intensity levels to compare the association between the origin and terminating factors. The area of each colored ribbon depicts the frequency of the samples related with the particular typing and pathogenic intensity expression

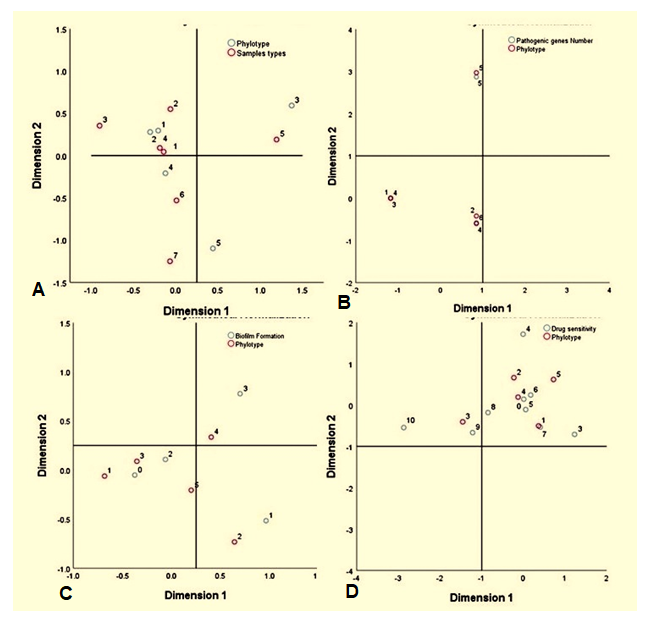

**FIGURE 9|** Correspondance analyses (CA) for the catégorial variables. Sample types, pathogenicity and phylogroup that are similar where this two-dimensional representation explain 100% of the total variation, with 68.55% explained by 1st dimension and 31.45% by the 2nd dimension. On the other hand, for the drug sensitivity and biofilm formation representation explains 87.25% variation by the 1st dimension and 12.75% by the 2nd dimension. Here A) Represents phylotypes (white circle) Vs sample types relation (red circle); B) Represents phylotypes (red circle) Vs pathogenic gees relation (white circle); C) Represents phylotypes (red circle) Vs biofilm formation (white circle); D) Represents phylotypes (red circle) Vs drug sensitivity (white circle).
