## Supplementary Figure Legends for "Multidrug-Resistant Avian Pathogenic *Escherichia coli* Strains and Association of Their Virulence Genes in Bangladesh"

**Supplementary Figures**

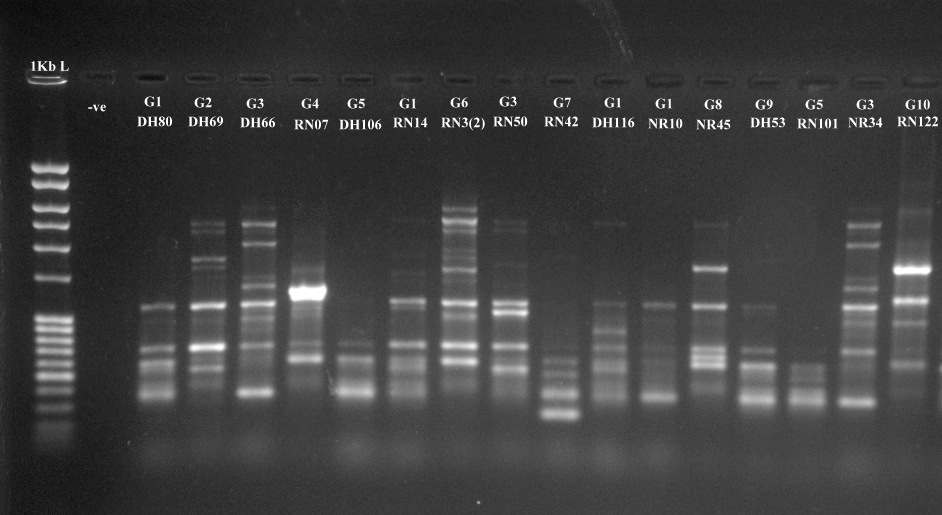

SUPPLEMENTARY FIGURE 1| RAPD patterns of bacterial isolate using primer 1283. Lane 2 is negative blank control and lanes1 is molecular ladders. Lanes 3–18 are samples DH80, DH69, DH66, RN07, DH106, RN14, RN3 (2), RN50, RN42, DH116, NR10, NR45, DH53, RN101, NR34, RN122 respectively representing group 1-10.

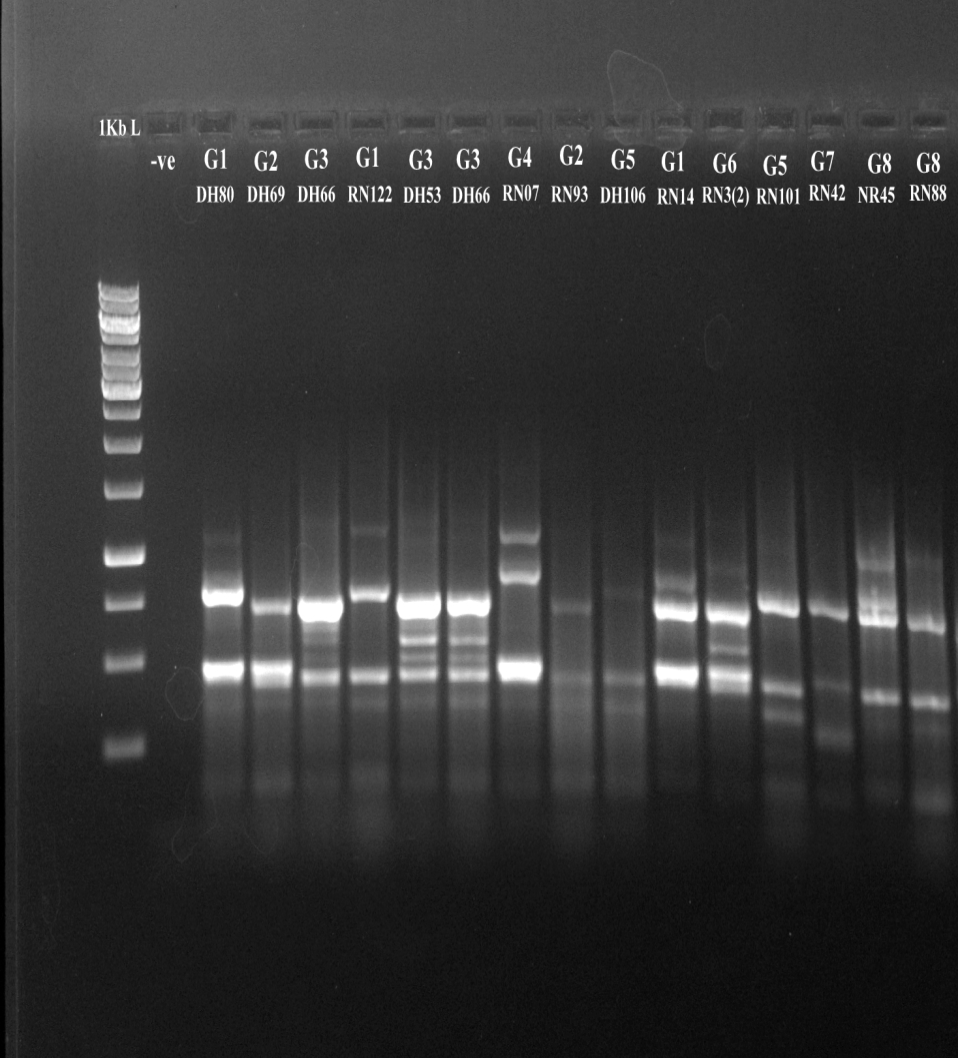

SUPPLEMENTARY FIGURE 2| ERIC-PCR patterns of bacterial isolate using primer ERIC1 and ERIC2. Lane 2 is negative blank control and lanes 1 is molecular ladders. Lanes 3–17 are samples DH80, DH69,DH66, RN122, DH53, DH66, RN07, RN93, DH106, RN14, RN3 (2), RN101, RN42, NR45, RN88 respectively representing group 1-8.

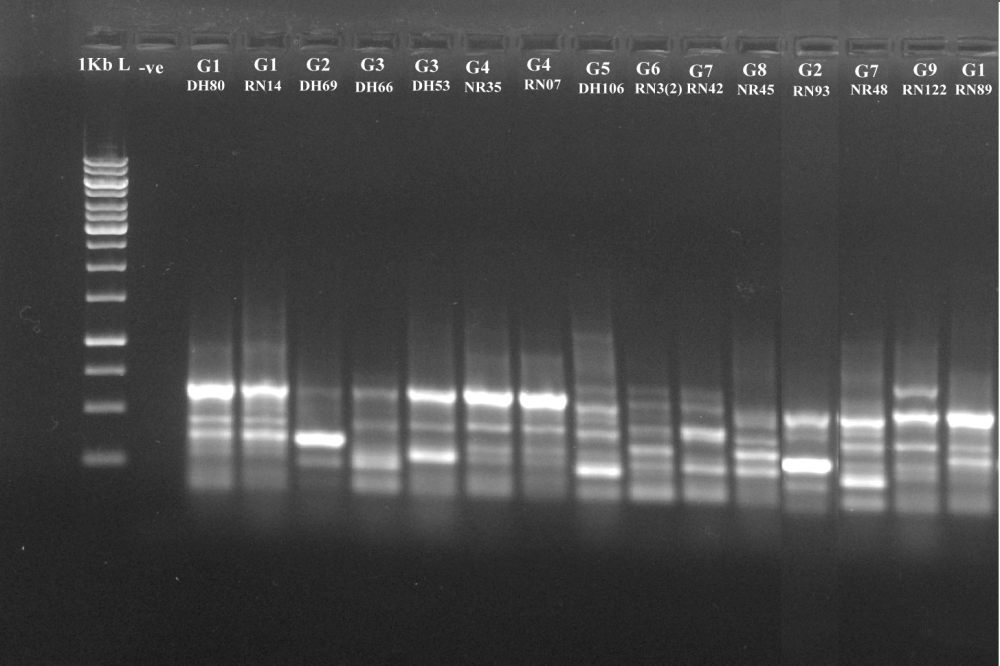

SUPPLEMENTARY FIGURE 3| BOX-PCR patterns of bacterial isolate using primer BOXA1R. Lane 2 is negative blank control and lanes 1 is molecular ladders. Lanes 3–17 are samples DH80, RN14, DH69, DH66, DH3, NR35, RN07, DH106, RN3 (2), RN42, NR45, RN93, NR48, RN122, RN89 respectively representing group 1-9.

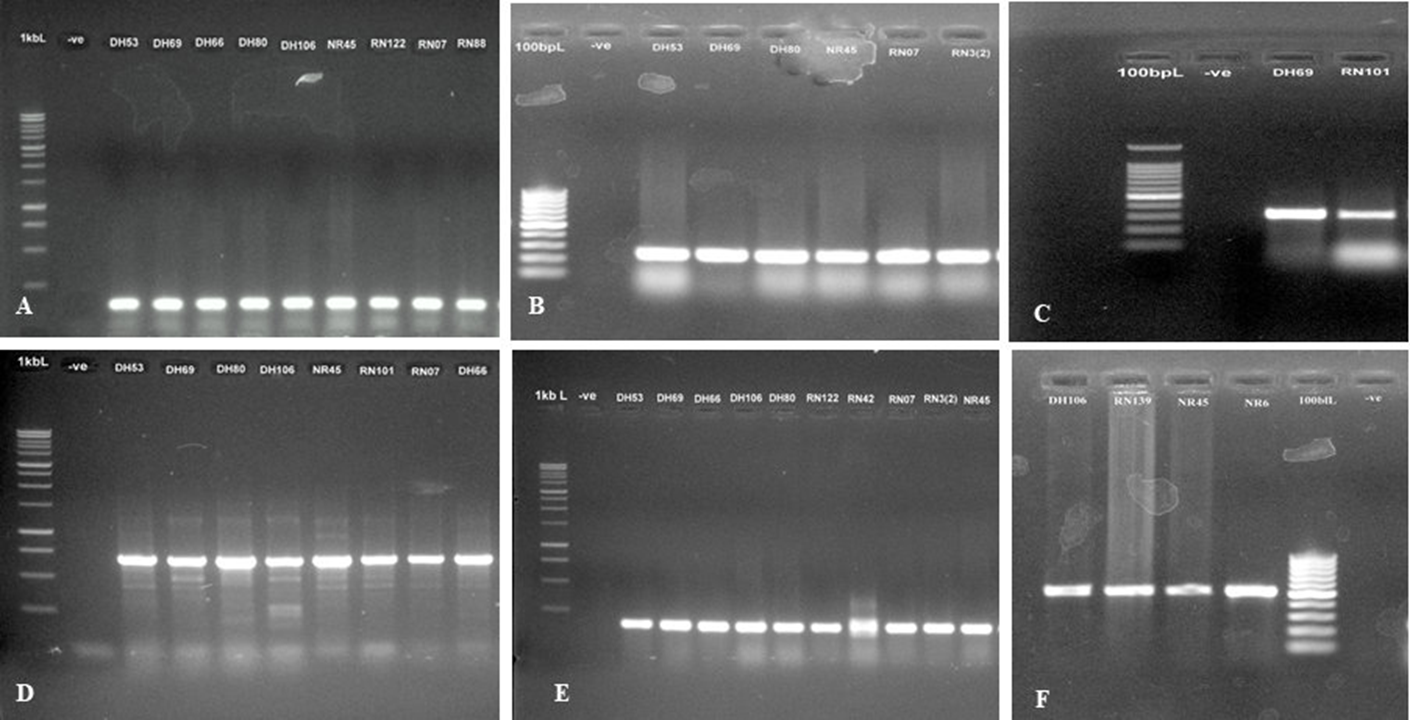

SUPPLEMENTARY FIGURE 4| PCR results for the detection of *E. coli* VG among the colibacillosis cases of Bangladeshi poultry samples. A) (*uidA*: 147bp) Lane 2 is negative blank control and lanes 1 is molecular ladders (1kb). Lanes 3-11 are strain DH53, DH69, DH66, DH80, DH106, NR45, RN122, RN07, RN88. B) (*crl*: 250bp) Lane 2 is negative blank control and lanes 1 is molecular ladders (100bp). Lanes 3–10 are strain DH53, DH69, DH80, NR45, RN07, RN3 (2). C) (*papC*: 328bp) Lane 2 is negative blank control and lanes 1 is molecular ladders (100bp). Lanes 3–4 are strain DH69, RN101. D) (*ial*: 650bp) Lane 2 is negative blank control and lanes 1 is molecular ladders (1kb). Lanes 3-10 are strain DH53, DH69, DH80, DH106, NR45, RN101, RN07, DH66.E) (*fimH*: 164bp) Lane 2 is negative blank control and lanes 1 is molecular ladders (1kb). Lanes 3-12 are strain DH53, DH69, DH66, DH106, DH80, RN122, RN42, RN07, RN3 (2), NR45.F) (*cjrC*: 518bp) Lane 6 is negative blank control and lanes 5 is molecular ladders (100bp). Lanes 1-4 are strain DH106, RN83, NR45, NR6.

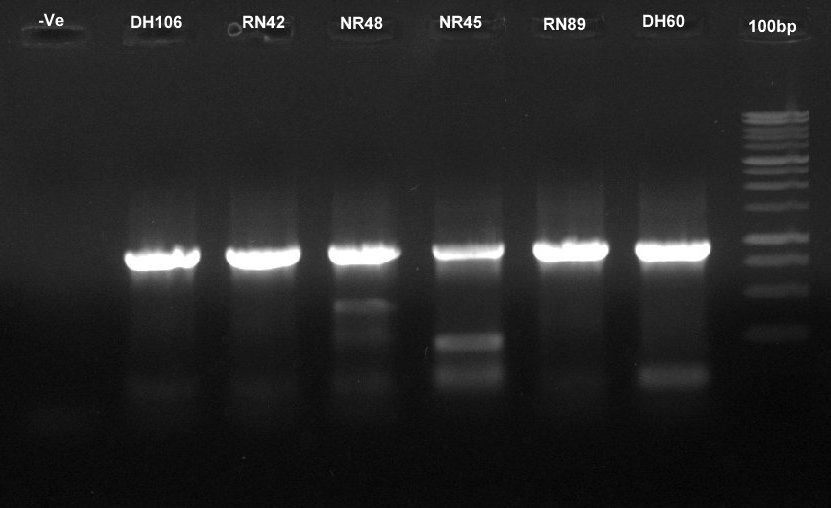

SUPPLEMENTARY FIGURE 5 | Representative PCR results for the detection of *Escherichia coli* phylotypes among the colibacillosis cases of Bangladeshi poultry samples. (*arp*A: 400bp), Here, Lane 8 is molecular ladders (100bp), Lane 1 is negative blank control and Lanes 2–7 are the strains DH106, RN42, NR48, NR45, RN89, DH60, respectively.
